# Whole-Brain Convergence of Real-World Visual Expertise beyond Faces: Meta-Analytic Evidence for Core–Adaptive Neural Architecture

**DOI:** 10.64898/2025.12.29.696823

**Authors:** Weilu Chai, Yuxin Bai, Xuemei Xie, Jimin Liang, Xiaoqi Huang, Janniko R. Georgiadis, Minghao Dong

**Author notes:** **Correspondence:** Professor Minghao Dong.

## Abstract

Visual expertise—the ability to discriminate highly similar exemplars quickly and accurately within a category—supports skilled performance across real-world domains and is supported by distributed neural systems. We focus on non-face expertise to test cross-domain convergence in acquired real-world visual skills, treating faces separately because socially embedded, sensitive-period-constrained processing could blur this inference. It remains unresolved whether non-face expertise across heterogeneous domains converges on a shared, domain-general whole-brain architecture, or instead recruits domain-contingent neural configurations that vary with task and stimulus demands. We conducted a coordinate-based meta-analysis of 22 task-fMRI studies spanning 11 real-world non-face expertise domains (579 participants, 210 peak-activation foci). Primary analysis revealed a robust, right-lateralized parieto-temporo-occipital circuit centered on the middle occipital gyrus, middle temporal gyrus, angular gyrus and adjoining inferior parietal lobule. We propose that this circuit constitutes a domain-general neural core that integrates fine-grained visual features with semantic associations while supporting attention-guided recognition of visually similar objects in expert performance. Subgroup and meta-regression analyses uncovered a complementary adaptive component, the engagement of which varied systematically with representational and contextual factors. Pictorial stimuli and expert–novice contrasts reliably strengthened recruitment of the right-hemisphere core, whereas symbolic stimuli engagement toward left temporal regions while selectively re-engaging right-core nodes. In addition, male-skewed samples showed attenuated left-hemisphere activation. Together, these findings delineate a stable right-hemisphere neural scaffold underlying non-face visual expertise, flexibly supplemented by left-hemisphere systems as a function of stimulus format, task demands, and demographic context, providing a whole-brain reference framework for future studies.

**Highlights:**

1. A coordinate-based meta-analysis integrates 22 whole-brain task-fMRI studies of real-world non-face visual expertise.
2. Results delineate a core–adaptive architecture supporting non-face expertise across domains.
3. A right-lateralized parieto–temporo–occipital core (AG, MOG, MTG, adjoining IPL) shows convergent recruitment across non-face domains.
4. Adaptive left-hemisphere regions (MTG/STG/ITG/IFG; plus FG/IOG/insula) are flexibly engaged with task and stimulus demands.
5. Findings refine neurocognitive accounts of perceptual expertise and provide a domain-spanning whole-brain reference framework for future studies.

**Graphical abstract:** 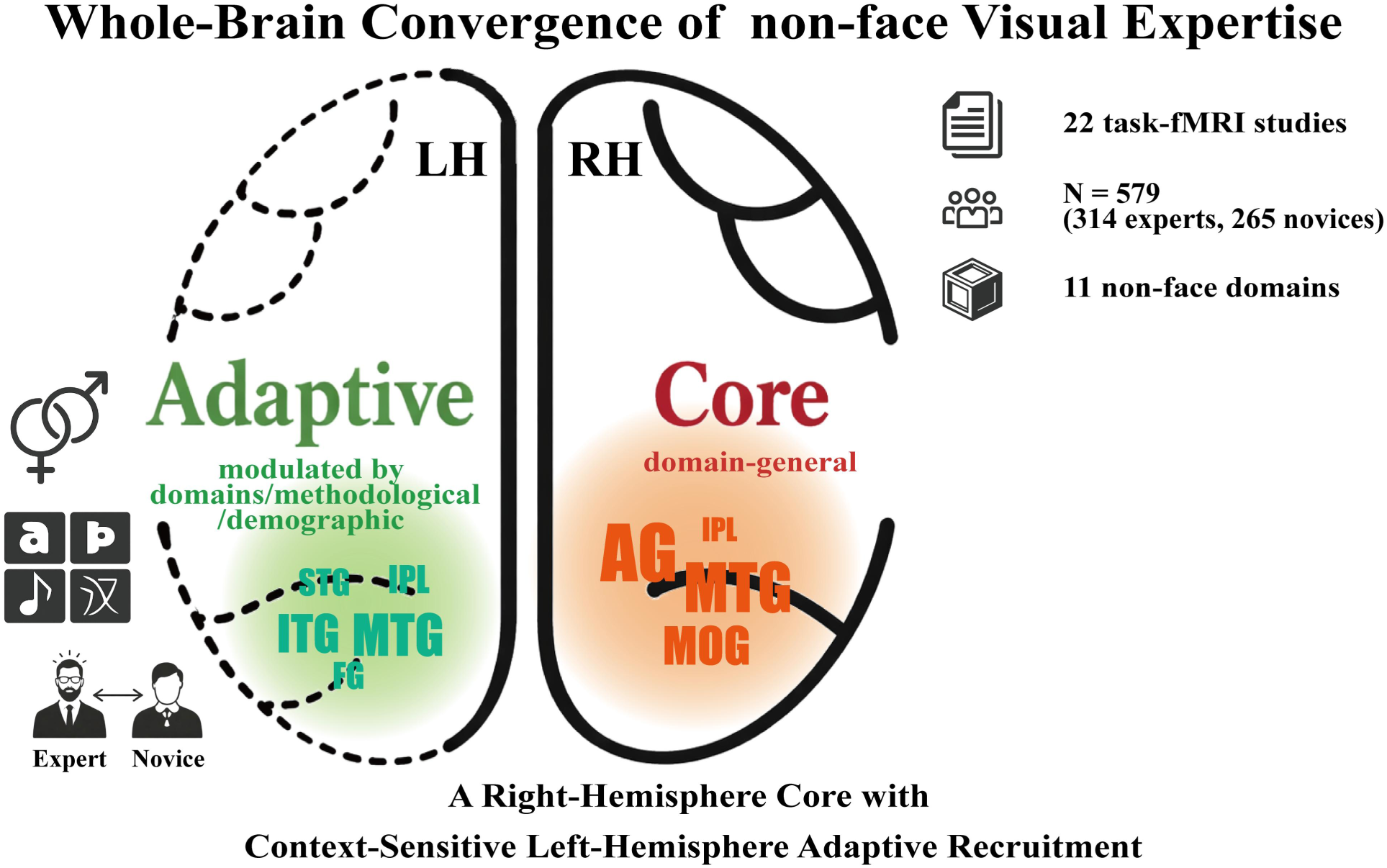

## 1. Introduction

Early theories of perceptual learning suggest that experience sharpens discrimination through changes in perceptual representations, over and above explicit rule-based strategies (Gibson, 1969). Visual expertise can be viewed as a real-world expression of this principle—a domain-specific capacity to reliably discriminate among highly similar exemplars within a visually homogeneous category (Harel et al., 2013), for example, distinguishing a Goldador from a Labrador. It underpins elite performance in a wide range of real-world activities, including word and music notation reading (Li et al., 2006; Wong and Gauthier, 2010), chess play (Bilalić et al., 2010), radiologic image interpretation (Melo et al., 2011), and fine-grained identification of vehicles, birds, organisms, minerals, and musical instruments (Gilaie-Dotan et al., 2012; Hoenig et al., 2011; Lee, and Kwon, 2011; Martens et al., 2018), as well as everyday domains such as face recognition (Young and Burton, 2018). At a behavioral level, visual expertise exhibits a convergent performance signature across domains, namely superior subordinate-level discrimination relative to non-experts, which typically emerges through years of prolonged, deliberate and domain-specific practice (Tanaka et al., 2005; Wu et al., 2024).

Over decades, expertise studies across diverse domains have progressively sharpened our understanding of visual expertise and the behavioral mechanisms that support it. Chess research provided an early perceptual–cognitive account of expertise, showing that experts’ advantage reflects rapid recognition of learned configurations rather than deeper search (Chase and Simon, 1973; Groot, 1965), and formalizing chunking and template models in which structured representations support fast encoding and anticipation (Chase and Simon, 1973; Ericsson and Kintsch, 1995; Gobet and Simon, 1996). This tradition also established methodological conventions that generalize across domains, including expert–novice contrasts with domain-relevant stimuli and intact-versus-randomized structure controls (Chase and Simon, 1973; Gobet and Simon, 1996). In radiology, early eye-movement studies showed that increasing experience reshapes scan paths and improves search efficiency during image interpretation (Kundel and La Follette Jr, 1972; Thomas and Lansdown, 1963); later accounts formalized this as a two-stage process in which an early screening stage flags candidate features for attention before explicit evaluation stage (Swensson, 1980). Subsequent gaze-tracking studies further demonstrated that a rapid holistic or global analysis can localize candidate abnormalities very early and guide subsequent focal verification (Kundel et al., 2007). Additionally, face recognition studies provided a benchmark for expertise at the level of individuation by highlighting holistic/configural processing—identity judgments depend on relations among features and whole-context encoding rather than independent parts (Tanaka and Farah, 1993; Young et al., 1987). Evidence from symbolic domains offered a parallel “global-to-local” pattern: word-reading models showed that recognition is strongly shaped by context and can be described as interactive, hierarchical processing across feature–letter–word levels (McClelland and Rumelhart, 1981; Reicher, 1969), and music-reading studies indicate that skilled readers rely on preview-based sampling to support fluent parsing beyond note-by-note scanning (Kinsler and Carpenter, 1995; Puurtinen, 2018). Taken together, these traditions converge on a domain-general behavioral hallmark of visual expertise, namely experience-driven sensitivity to diagnostic relational structure that enables rapid mapping from perceptual input to domain-relevant meaning, supported by efficient global-to-focal processing. This convergence naturally raises the question of whether a corresponding unified neural architecture underlies visual expertise across domains.

Neural plasticity refers to the nervous system’s capacity to reorganize its structure and function, with experience-dependent change being a central mechanism in learning and expertise (Pascual-Leone et al., 2005; Zatorre et al., 2012). Within vision, perceptual learning provides a tractable model of plasticity in the mature visual system: repeated practice yields durable improvements in detection or fine discrimination, accompanied by experience-dependent reweighting within visual cortical circuits and task-dependent top-down influences on visual processing (Sasaki et al., 2010; Watanabe and Sasaki, 2015). Long-term deliberate practice in real-world domains can be viewed as an ecological extension of these principles, motivating extensive neuroimaging work on how sustained experience and training sculpt the neural representations that support visual expertise. Early work identified a fusiform face area (FFA) showing selective responses during face recognition (Kanwisher et al., 1997), and subsequent evidence suggested that FFA responses can also increase with acquired visual expertise following training on novel objects (Gauthier et al., 1999). However, subsequent research across diverse expertise domains has more often implicated distributed patterns of expertise-related activation extending beyond ventral temporal cortex (Harel, 2016; Harel et al., 2013). For example, taxonomic biologists recruit the right middle temporal gyrus (MTG) and inferior parietal lobule (IPL) when discerning subtle morphological regularities (Lee, and Kwon, 2011). Musicians reading notation recruit bilateral ventral temporal cortex together with left-lateralized perisylvian regions (e.g., left inferior frontal gyrus (IFG) and supramarginal regions) and intraparietal cortex, consistent with symbol decoding and learned mappings from visual notation to domain-relevant representations (Mongelli et al., 2017). In car experts, car-selective activity is widespread, extending beyond early visual cortex into far-peripheral visual representations and face/object-selective regions, and additionally recruiting regions outside occipitotemporal cortex (e.g., precuneus, intraparietal sulcus, and prefrontal cortex) (Harel et al., 2010). Chess masters recruit bilateral parieto-temporo-occipital junctions and retrosplenial cortex, together with left supramarginal gyrus, during recognition of chess pieces and their spatial configurations (Bilalić et al., 2012). Across these domains, expertise effects recur in a parieto-temporo-occipital set of regions implicated in linking visual structure to learned associations, while additional activations vary with representational format and task demands. These observations motivate a cross-domain neural hypothesis of visual expertise: non-face real-world visual expertise is supported by a relatively stable parieto-temporo-occipital core that integrates high-precision visual information with semantic associations, complemented by adaptive extensions whose engagement varies systematically with stimulus format, representational demands, and task context. Crucially, this account predicts both convergence across expertise domains and principled heterogeneity across experimental paradigms, predictions that cannot be adequately evaluated within individual, domain-bound neuroimaging studies.

At the same time, cross-domain synthesis faces an important complication: visual expertise varies across domains in the perceptual and interpretive demands it imposes. In particular, faces are a distinctive—and potentially confounding—case for cross-domain inference. Face recognition is intrinsically tied to social identity and interpersonal meaning, and reliably engages processes that extend beyond perceptual analysis (Haxby et al., 2000; Young and Burton, 2018). Moreover, face processing is shaped by early developmental constraints, with evidence from early visual deprivation indicating that typical holistic face processing depends on experience during a sensitive period in infancy (Gauthier and Nelson, 2001; Grand et al., 2004; McKone et al., 2007). By contrast, non-face visual expertise is typically acquired later in life, embedded in human–object interaction rather than social identity recognition, and is not subject to the same infancy-limited constraint (Harel et al., 2013; McKone et al., 2007; Wu et al., 2024). Including faces would therefore risk conflating domain-general mechanisms of visual expertise with socially and developmentally specific processes unique to face recognition. Accordingly, while face processing has been extensively synthesized through multiple coordinate-based meta-analyses (Bzdok et al., 2011; Dricu and Frühholz, 2016; Müller et al., 2018b; Zinchenko et al., 2018), non-face expertise has largely been examined within single domains (e.g., musical expertise (Criscuolo et al., 2022) or skilled word reading (Murphy et al., 2019)), limiting direct evaluations of whether different non-face domains converge on a shared whole-brain architecture. Addressing this gap is theoretically critical for distinguishing domain-general neural mechanisms from task- or format-dependent adaptations, thereby providing convergent evidence bearing on the proposed core–adaptive account.

A further challenge for cross-domain synthesis is that non-face expertise paradigms differ in representational format and in how expertise is operationalized. Accordingly, heterogeneity is expected to arise from at least three psychologically grounded sources. First, stimulus format matters: symbolic stimuli such as words or notation are composed of simple visual primitives governed by combinatorial rules, whereas pictorial stimuli such as objects or scenes are defined by rich feature conjunctions (Grill-Spector and Weiner, 2014; Koelsch et al., 2004; Patel, 2003). Second, experimental contrasts differ: some studies rely on expert–novice comparisons within a domain, whereas others contrast domain-relevant and irrelevant categories within experts, each approach introducing distinct interpretive trade-offs (Haller and Radue, 2005; Lee, and Kwon, 2011)). Third, demographic factors, including sex and age, may further modulate hemispheric organization and neural selectivity (Carp et al., 2011; Vanston and Strother, 2017). These dimensions render the core–adaptive account explicitly testable at the whole-brain level and motivate prespecified subgroup and meta-regression analyses.

In the present study, we evaluate the core–adaptive account through a cross-domain, whole-brain coordinate-based synthesis of task-fMRI studies of non-face real-world visual expertise. We first quantify convergence across all included domains to assess whether a shared expertise-related architecture can be identified. We then examine systematic modulation by stimulus format and experimental contrast using subgroup analyses and assess demographic influences via meta-regression. Together, these analyses provide a quantitative assessment of the core–adaptive account by distinguishing domain-general neural convergence from principled, demand-dependent variation across real-world expertise domains.

## 2. Methods

### 2.1. Literature Search Strategy

We conducted a coordinate-based meta-analysis following established practices in prior research, in accordance with standard neuroimaging meta-analytic guidelines and the Preferred Reporting Items for Systematic Reviews and Meta-Analyses (PRISMA) statement (Müller et al., 2018a; Page et al., 2021; Xiao et al., 2024). The protocol was pre-registered on the Open Science Framework (https://osf.io/dpcy5) to ensure methodological transparency and reproducibility.

A comprehensive search was conducted for peer-reviewed fMRI studies examining visual expertise across diverse non-face object domains. All English-language publications published between March 2001 and March 2025 were considered, with no restrictions on geographic origin. Literature was retrieved from PubMed, PsycINFO, and Web of Science using combinations of the following keywords: (“fMRI”) AND (“expert” OR “expertise” OR “professional” OR “training” OR “train” OR “prodigy”) AND (“visual” OR “vision”), excluding terms related to non-human animals or clinical populations (e.g., “monkey,” “mice,” “animal,” “patient”). Detailed search formulas for each database are provided in *Supplementary Table 1*.

### 2.2. Selection Criteria

The systematic review process is summarized in the PRISMA flow diagram (*Figure 1*), with the full checklist provided in *Supplementary Table 4*. The search yielded 1254 non-duplicate articles, including 1223 from database queries and 31 from manual searches, all managed using EndNote. Titles and abstracts were screened according to the following predefined inclusion criteria:

1. **Article type**: Only peer-reviewed research articles published in English were included; dissertations, theses, review articles, and commentaries were excluded.
2. **Participant characteristics**: Studies were required to involve healthy adult participants (aged 18–65 years). Studies involving children, older adults (>65 years), clinical populations, or non-human subjects were excluded.
3. **Experimental paradigm**: Only fMRI studies employing controlled tasks involving active visual recognition (e.g., recognition, matching, detection) were included. Resting-state fMRI studies were excluded.
4. **Domain scope**: Studies of visual expertise in non-face object domains (e.g., words, musical notation, radiographs) were included; face-related studies were excluded.
5. **Expertise Validation**: Studies must prospectively validate expert status through either objectively quantified domain-specific performance metrics or substantiated professional training/experience.
6. **Expertise Effect Isolation**: Included studies were required to isolate expertise-related neural responses by controlling for non-expertise-related cognitive processes (e.g., expert-novice contrasts or cross-domain designs).
7. **Neuroimaging Methodology**: Studies were required to report task-based whole-brain voxel-wise activation results with peak coordinates in standard space (Talairach or MNI). Studies focusing solely on connectivity analyses or ROI-based approaches were excluded.

**Figure 1.**
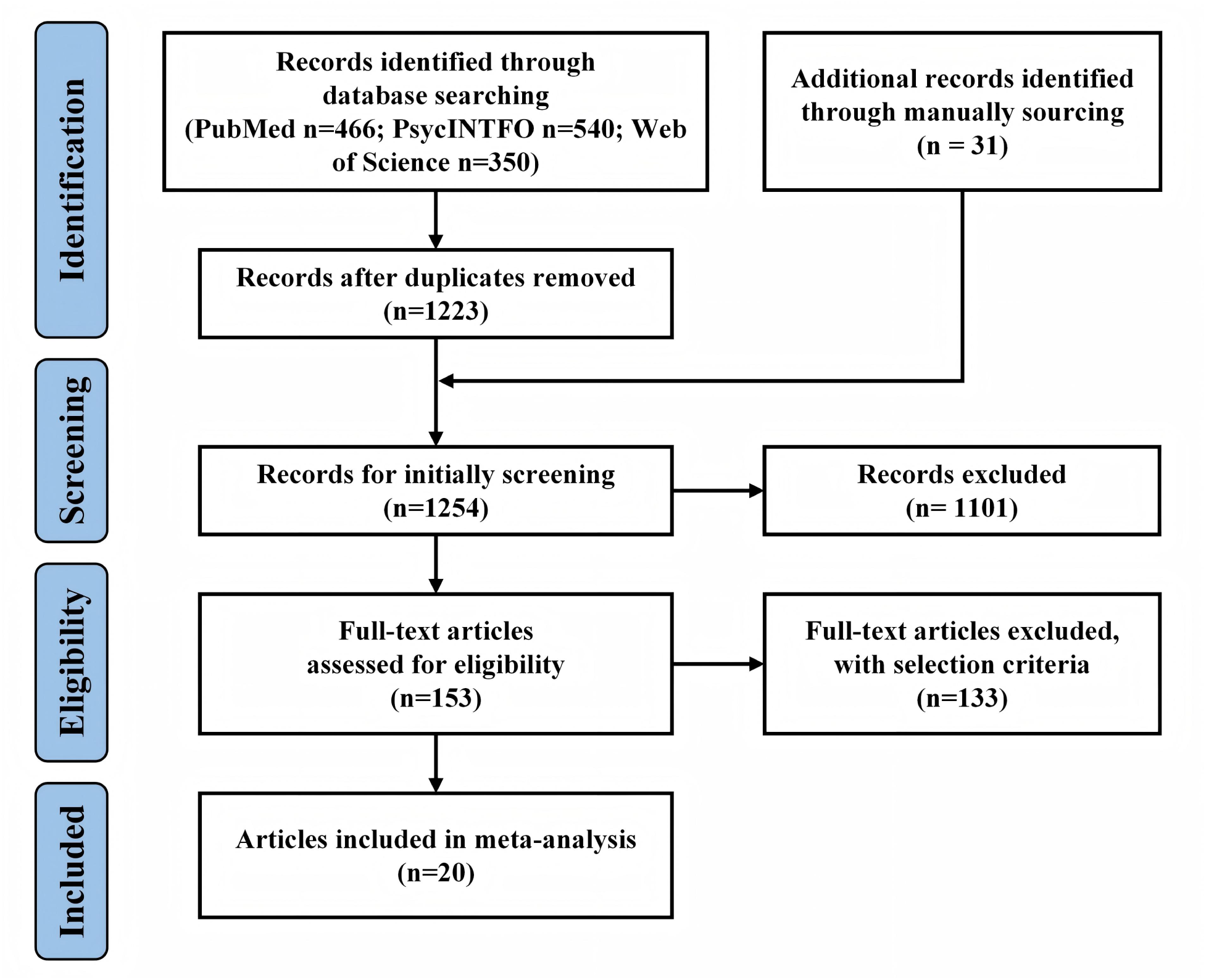
The diagram of the systematic review search process.

Initial screening excluded 1101 articles due to being review articles, involving non-human or clinical populations, targeting non-visual domains, or lacking fMRI methodology. A total of 153 studies underwent full-text assessed for eligibility, resulting in 22 articles meeting all inclusion criteria. The remaining 131 were excluded due to issues related to task paradigm, expertise validation, confound control, or imaging methodology. In addition, two more studies were excluded due to missing peak coordinates, and author inquiries were unsuccessful (Harel et al., 2010; McGugin et al., 2015). The final sample comprised 22 studies from 20 articles, including two articles each reporting two distinct experiments.

### 2.3. Data Extraction

For each of the 22 included studies, methodological information was extracted, including the first author, publication year, expertise domain, stimulus type, expertise validation method, task paradigm, experimental contrast, sample size, expert age (mean), expert male ratio, and t-value threshold (see *Table 1* for details). For meta-analytic preprocessing, t-value thresholds were recorded across all studies and, when only p-values were reported, converted using the SDM-provided conversion tool (https://www.sdmproject.com). The primary neuroimaging data consisted of peak activation coordinates reported in standard space (Talairach or MNI), along with their corresponding statistical values (p-values or z-values). In cases where the magnitude was not directly available, activations were coded as positive (“p”) or negative (“n”) in accordance with Seed-based d Mapping with Permutation of Subject Images (SDM-PSI) conventions (Albajes-Eizagirre et al., 2019).

**Table 1.** Methodological details of 22 studies (from 20 included articles) with the visual expertise effect included in the meta-analysis.

| Author name | Year | Expertise domain | Stimulus types | Expertise validation method | Task paradigm | Experimental contrast | Sample size |  | VE age (mean) | VE man ratio | t-value threshold |
| --- | --- | --- | --- | --- | --- | --- | --- | --- | --- | --- | --- |
|  |  |  |  |  |  |  | VE* | non-VE* |  |  |  |
| Daniele Schön et al. (Schön et al., 2002) | 2002 | musical notation | symbolic | > 12 y | Object recognition | Intra-group comparison: musical notation > verbal or number notation | 9 | NA | 37** | 0.56 | 1.75 |
| Geng Li et al. (Li et al., 2006) | 2006 | Chinese word | symbolic | native speakers | Silent word recognition | Inter-group comparison | 12 | 12 | 67 | 0.5 | 4.02 |
| Campitelli et al. (Campitelli et al., 2007) | 2007 | Chess | pictorial | Elo*: avg. 1978 | Matching recognition | Intra-group comparison: chess game > geometric shapes/dots | 5 | NA | 24.6 | NA | 1.86 |
| Bilalic et al. (Bilalić et al., 2010) | 2010 | Chess | pictorial | Elo: avg. 2108 | Pattern recognition | Inter-group comparison | 8 | 15 | 30 | 1 | 1.72 |
| Lee et al. (Lee, and Kwon, 2011) | 2011 | Organism | pictorial | doctorates | Pattern recognition | Inter-group comparison | 8 | 8 | 37.9 | 1 | 1.76 |
| Hoenig et al. (Hoenig et al., 2011) | 2011 | Musical instruments | pictorial | mean 27y | Object recognition | Inter-group comparison | 20 | 20 | 37 | 0.7 | 1.69 |
| Marcin Szwed et al. (Szwed et al., 2014) | 2014 | Chinese word | symbolic | native speakers | Easy oddball detection | Inter-group comparison | 11 | 14 | 25** | 0.36 | 1.71 |
|  |  | French word | symbolic | native speakers | Easy oddball detection | Inter-group comparison | 14 | 11 | 25.5** | 0.36 | 1.71 |
| Valeria Mongelli et al. (Mongelli et al., 2017) | 2017 | Musical notation | symbolic | mean 5.7y | 1-back matching | Inter-group comparison | 21 | 23 | 32.6 | 0.57 | 1.68 |
| Farah Martens et al. (Martens et al., 2018) | 2018 | Bird | pictorial | mean 8.6y | 1-back matching | Inter-group comparison | 20 | 37 | 26.4 | 0.75 | 1.67 |
|  |  | Mineral | pictorial | Mean 6.4y | 1-back matching | Inter-group comparison | 17 | 40 | 25.3 | 0.59 | 3.25 |
| Wanwan Guo et al. (Guo et al., 2022) | 2022 | Chinese words | symbolic | native speakers | Chinese word recognition | Intra-group comparison: real words > pseudowords | 51 | NA | 23.4 | 0.49 | 1.66 |
| Wong et al. (Wong and Gauthier, 2010) | 2010 | Musical notation | symbolic | mean 14.4y | 1-back matching | Inter-group comparison | 10 | 10 | 21.4 | 0.2 | 1.73 |
| Krawczyk et al. (Krawczyk et al., 2011) | 2011 | Chess | pictorial | mean 16y | 1-back matching | Inter-group comparison | 6 | 6 | 23 | 1 | 4.14 |
| Bilalić et al. (Bilalić et al., 2011) | 2011 | Chess | pictorial | Elo: avg. 2130 | Pattern recognition | Inter-group comparison | 8 | 8 | 29 | 1 | 3.79 |
| Wu et al. (Wu et al., 2012) | 2012 | Chinese word | symbolic | native speakers | Word recognition | Inter-group comparison | 11 | 11 | 43.9 | 0.55 | 2.53 |
| Bilalić et al. (Bilalić et al., 2012) | 2012 | Chess | pictorial | Elo: avg. 2108 | Pattern recognition | Inter-group comparison | 8 | 15 | 30 | 1 | 1.72 |
| Bartlett et al. (Bartlett et al., 2013) | 2013 | Chess | pictorial | mean 16y | 1-back matching | Inter-group comparison | 11 | 11 | 23 | 1 | 3.55 |
| Sharon et al. (Gilaie-Dotan et al., 2012) | 2012 | Cars | pictorial | Recognition rate >83% | 1-back matching | Inter-group comparison | 12 | 9 | 23.4 | 1 | 1.73 |
| Sven et al. (Haller and Radue, 2005) | 2005 | radiologic images | pictorial | mean 6.9y | Object recognition | Intra-group comparison: radiologic images > electron microscopic images | 12 | NA | 35.8 | 0.83 | 3.5 |
| Marcio et al. (Melo et al., 2011) | 2011 | radiologic images | pictorial | mean 11.6y | Object recognition | Intra-group comparison: CXR* with lesions > CXR with animals or letter | 25 | NA | 35.9 | 0.64 | 1.68 |
| Duan et al. (Duan et al., 2012) | 2012 | Chinese Chess | pictorial | mean 13.67y | Pattern recognition | Inter-group comparison | 15 | 15 | 28.7 | 0.67 | 1.7 |

**Note:** Items marked with \* denote abbreviations: Elo refers to the international chess Elo rating scale; CXR refers to chest X-ray images; VE refers to visual experts in their domain of visual expertise; non-VE refers to non-experts in the visual expertise domain of VE. Items marked with \*\* indicate studies where only an age range (e.g., 24-50 years) is provided. In these cases, the midpoint of the range (37 years) was used as the representative value.

All inclusion decisions and data extractions were independently performed by two researchers (Weilu Chai and Minghao Dong), with discrepancies resolved through discussion until full agreement was reached.

### 2.4. Coordinate-based Meta-analysis

All coordinate-based meta-analyses—including a primary meta-analysis, subgroup analyses, and meta-regression analyses—were conducted using SDM-PSI version 6.22 (Albajes-Eizagirre et al., 2019). This approach integrates standard voxel-wise statistical tests with subject-based permutation procedures, thereby improving statistical power and minimizing the reliance on restrictive spatial assumptions that often limit prior coordinate-based methods (Albajes-Eizagirre et al., 2019). The primary meta-analysis aimed to identify consistent brain activation patterns associated with visual expertise across all included studies. Subgroup analyses grouped studies based on discrete potential modulatory factors, while meta-regression analyses tested continuous moderators within the primary meta-analytic model (see Section 2.6 for details).

Following official guidelines from the SDM project (www.sdmproject.com), we selected the voxel-based morphometry (VBM) gray matter modality. Preprocessing employed a 20 mm full width at half maximum (FWHM) kernel. Meta-analytic means were estimated using 50 multiple imputations, and 1,000 permutations were performed to enable family-wise error (FWE) correction. Statistical significance was determined using threshold-free cluster enhancement (TFCE), with results considered significant at *p_TFCE-corr_ < 0.05*. This thresholding method was chosen for its ability to balance sensitivity and specificity (Albajes-Eizagirre et al., 2019; Smith and Nichols, 2009). A 10-voxel extent threshold was applied to exclude spurious clusters. Final z-maps were visualized using MRIcron.

### 2.5. Heterogeneity, Sensitivity, Publication Bias, and Quality Assessment

Statistical heterogeneity across studies was assessed using the *I²* statistic, which quantifies the proportion of variability attributable to between-study differences rather than sampling error (Higgins and Thompson, 2002). Values of *I²* were interpreted as negligible (<10%), low (10% ∼ 39%), moderate (40% ∼ 59%), high (60% ∼ 89%) and very high (≥90%) heterogeneity (Schober et al., 2021). To evaluate the robustness of meta-analytic findings, jackknife sensitivity analyses were conducted within the SDM-PSI framework by sequentially excluding one study at a time (Radua and Mataix-Cols, 2009). Potential publication bias was assessed using Egger’s tests and visual inspection of funnel plot asymmetry (Egger et al., 1997). Quality assessment was independently performed for all included studies by two reviewers (Weilu Chai and Minghao Dong), with discrepancies resolved through discussion until consensus was reached.

### 2.6. Subgroup analysis and Meta-regression analysis

Subgroup and meta-regression analyses were conducted within the SDM-PSI framework to assess how the modulatory factors influenced expertise-related activation patterns.

Subgroup analyses were conducted based on two potential factors:

1. Stimulus type: symbolic (e.g., musical notation, written words) vs. pictorial (e.g., radiographs, chessboards). Studies were accordingly grouped into symbolic and pictorial stimuli subgroups.
2. Experimental contrast: inter-group comparisons (experts vs. novices) vs. intra-group comparisons (domain-relevant vs. irrelevant stimuli). Studies were categorized into inter-group and intra-group subgroups.

These groupings are reflected in the stimulus types and experimental contrast columns of *Table 1*, respectively.

Meta-regression analyses assessed the continuous influence of demographic covariates—proportion of male experts and mean age of experts—on voxel-wise activation across the primary meta-analytic space. Analyses proceeded in a stepwise fashion: first, male ratio was entered as a single regressor; next, mean age was added to the model to evaluate its incremental effect beyond variance explained by male ratio.

All subgroup and meta-regression results were thresholded at *p_TFCE-corr_ < 0.05* to ensure consistency with the primary analysis.

## 3. Results

### 3.1. Demographic and Study Characteristics

This coordinate-based meta-analysis included 210 peak-activation foci extracted from 22 task-based fMRI studies reported in 20 articles published between 2002 and 2022, involving 314 visual experts (mean age = 30.90 ± 10.09 years; 71.05% male) and 265 novices. The studies spanned 11 distinct expertise domains, including chess, music, radiology, and Chinese characters (see *Table 1* for full details).

In terms of stimulus type, 8 studies (36.36%) employed symbolic stimuli (e.g., musical notation, written language), while 14 studies (63.64%) used pictorial stimuli (e.g., X-ray images, natural objects, chessboards). Both musical notation and written language are socially learned symbolic systems that rely on abstract rule-based decoding and share close cognitive and neural substrates, and were therefore classified as symbolic stimuli in this study (Patel, 2003; Patel, 2007; Patel, 2012). For experimental contrast, 17 studies (77.27%) used inter-group designs, contrasting experts with novices on domain-relevant stimuli. The remaining 5 studies (22.73%) employed intra-group contrasts, comparing experts’ responses to domain-relevant versus irrelevant stimuli.

Following quality assessment, all included studies were rated as high quality. The assessment criteria and individual study scores are detailed in *Supplementary Tables 2 and 3*.

### 3.2. Primary Meta-Analysis Results

The primary meta-analysis across all 22 task-based fMRI studies identified two significant clusters of activation associated with visual expertise (*Table 2; Figure 2*). The largest cluster encompassed regions in the right middle occipital gyrus (MOG), MTG, angular gyrus (AG), and inferior parietal (excluding supramarginal and angular) lobule (IPL), with peak activation located in the right MOG (*MNI: 46, –64, 26; SDM-Z = 4.719; p_TFCE-corr_ <0.05; cluster size = 1414 voxels*). Another cluster was observed in the right MTG (*MNI: 62, –44, –6; SDM-Z = 3.705; p_TFCE-corr_ <0.05; cluster size = 219 voxels*).

**Figure 2.**
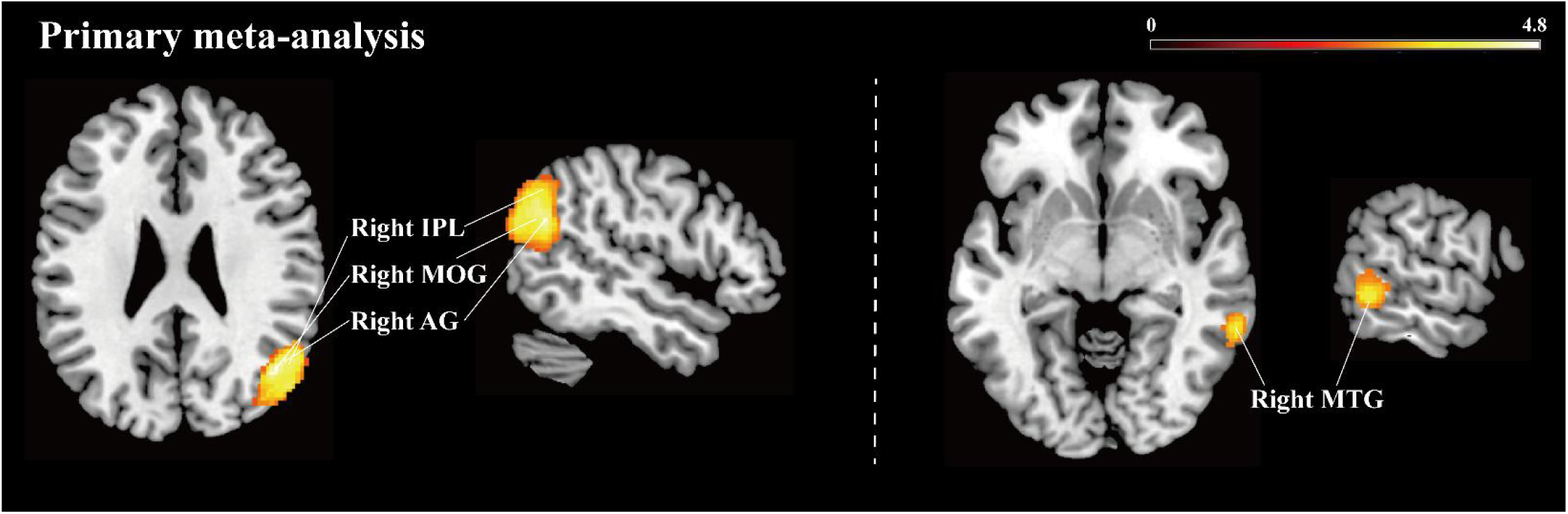
Z-map of significant activations results associated with the visual expertise effect identified by the primary meta-analysis at *p_TFCE-corr_* < 0.05. *Note: In heat map representations, a stronger yellow indicates a higher positive z value.

**Table 2.**
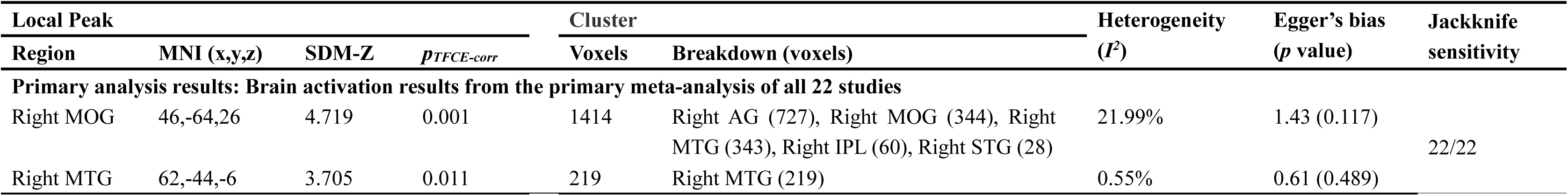
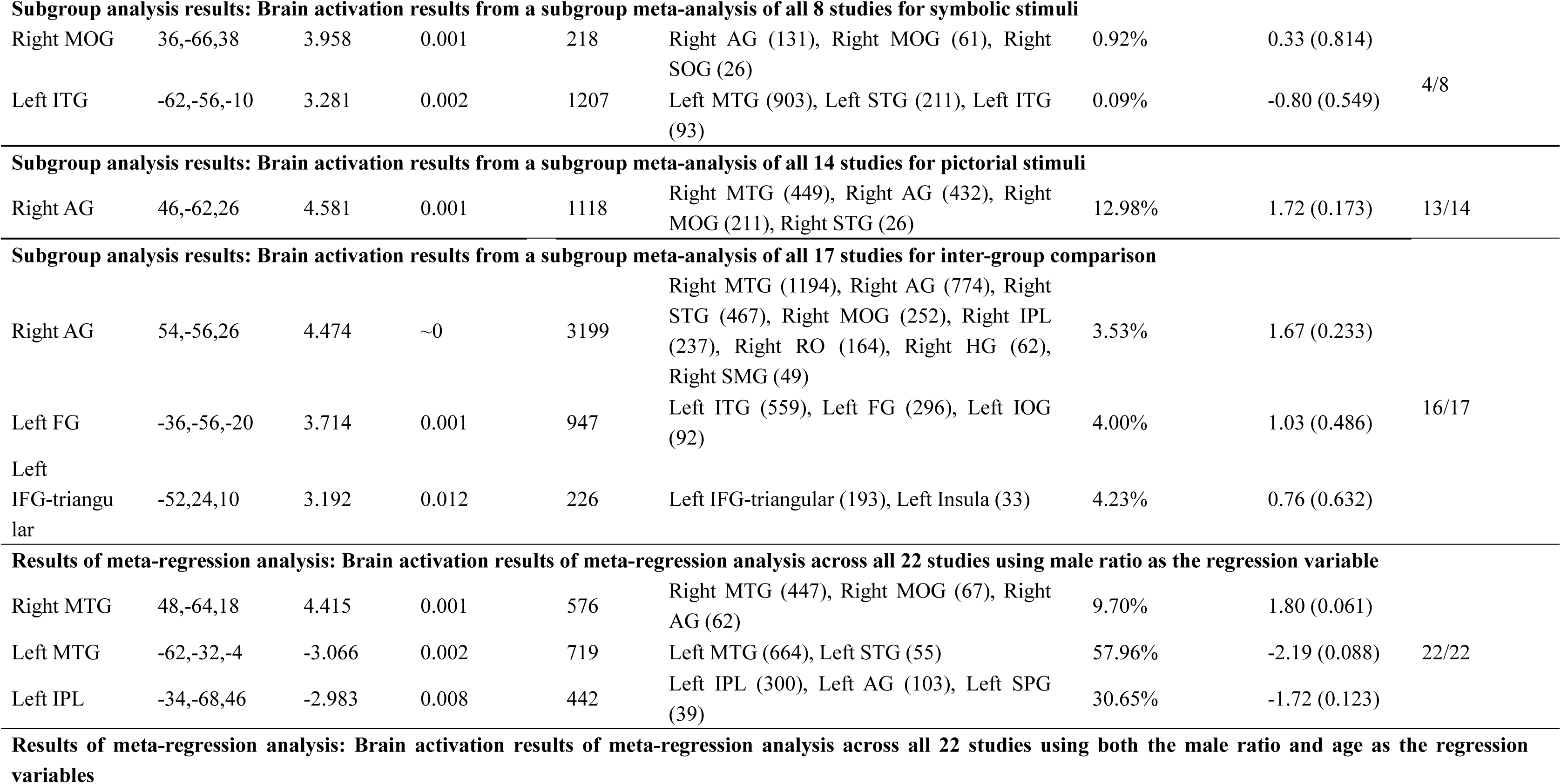
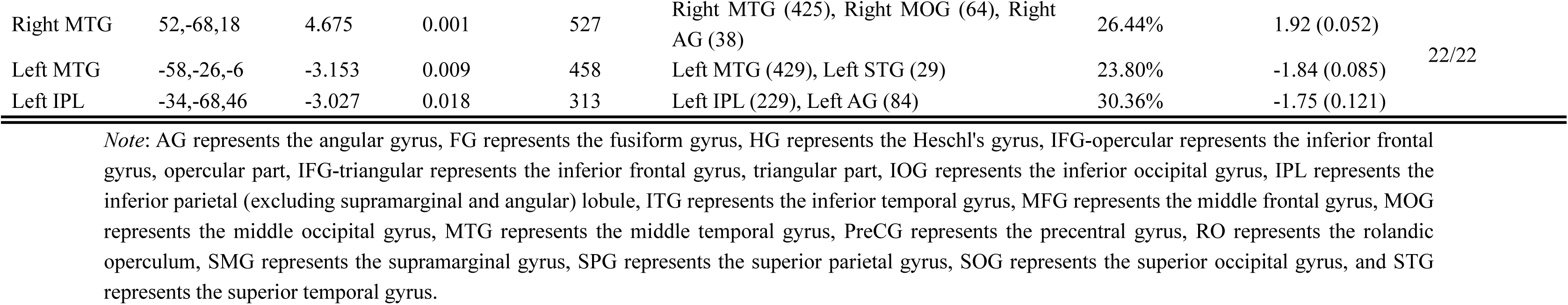
Peak coordinates of significant activations associated with the visual expertise effect identified by the meta-analysis at *p_TFCE-corr_* < 0.05.

| Local Peak |  |  |  | Cluster |  | Heterogeneity<br>(*I*<sup>2</sup>) | Egger's bias<br>(*p* value) | Jackknife<br>sensitivity |
| --- | --- | --- | --- | --- | --- | --- | --- | --- |
| Region | MNI (x,y,z) | SDM-Z | *p*<sub>TFCE-corr</sub> | Voxels | Breakdown (voxels) |  |  |  |
| Primary analysis results: Brain activation results from the primary meta-analysis of all 22 studies |  |  |  |  |  |  |  |  |
| Right MOG | 46,-64,26 | 4.719 | 0.001 | 1414 | Right AG (727), Right MOG (344), Right MTG (343), Right IPL (60), Right STG (28) | 21.99% | 1.43 (0.117) | 22/22 |
| Right MTG | 62,-44,-6 | 3.705 | 0.011 | 219 | Right MTG (219) | 0.55% | 0.61 (0.489) |  |
| **Subgroup analysis results: Brain activation results from a subgroup meta-analysis of all 8 studies for symbolic stimuli** |  |  |  |  |  |  |  |  |
| Right MOG | 36,-66,38 | 3.958 | 0.001 | 218 | Right AG (131), Right MOG (61), Right SOG (26) | 0.92% | 0.33 (0.814) | 4/8 |
| Left ITG | -62,-56,-10 | 3.281 | 0.002 | 1207 | Left MTG (903), Left STG (211), Left ITG (93) | 0.09% | -0.80 (0.549) |  |
| Right AG | 46,-62,26 | 4.581 | 0.001 | 1118 | Right MTG (449), Right AG (432), Right MOG (211), Right STG (26) | 12.98% | 1.72 (0.173) | 13/14 |
| Right AG | 54,-56,26 | 4.474 | ~0 | 3199 | Right MTG (1194), Right AG (774), Right STG (467), Right MOG (252), Right IPL (237), Right RO (164), Right HG (62), Right SMG (49) | 3.53% | 1.67 (0.233) | 16/17 |
| Left FG | -36,-56,-20 | 3.714 | 0.001 | 947 | Left ITG (559), Left FG (296), Left IOG (92) | 4.00% | 1.03 (0.486) |  |
| Left IFG-triangular | -52,24,10 | 3.192 | 0.012 | 226 | Left IFG-triangular (193), Left Insula (33) | 4.23% | 0.76 (0.632) |  |
| Right MTG | 48,-64,18 | 4.415 | 0.001 | 576 | Right MTG (447), Right MOG (67), Right AG (62) | 9.70% | 1.80 (0.061) | 22/22 |
| Left MTG | -62,-32,-4 | -3.066 | 0.002 | 719 | Left MTG (664), Left STG (55) | 57.96% | -2.19 (0.088) |  |
| Left IPL | -34,-68,46 | -2.983 | 0.008 | 442 | Left IPL (300), Left AG (103), Left SPG (39) | 30.65% | -1.72 (0.123) |  |
| Right MTG | 52,-68,18 | 4.675 | 0.001 | 527 | Right MTG (425), Right MOG (64), Right AG (38) | 26.44% | 1.92 (0.052) | 22/22 |
| Left MTG | -58,-26,-6 | -3.153 | 0.009 | 458 | Left MTG (429), Left STG (29) | 23.80% | -1.84 (0.085) |  |
| Left IPL | -34,-68,46 | -3.027 | 0.018 | 313 | Left IPL (229), Left AG (84) | 30.36% | -1.75 (0.121) |  |

*Note:* AG represents the angular gyrus, FG represents the fusiform gyrus, HG represents the Heschl's gyrus, IFG-opercular represents the inferior frontal gyrus, opercular part, IFG-triangular represents the inferior frontal gyrus, triangular part, IOG represents the inferior occipital gyrus, IPL represents the inferior parietal (excluding supramarginal and angular) lobule, ITG represents the inferior temporal gyrus, MFG represents the middle frontal gyrus, MOG represents the middle occipital gyrus, MTG represents the middle temporal gyrus, PreCG represents the precentral gyrus, RO represents the rolandic operculum, SMG represents the supramarginal gyrus, SPG represents the superior parietal gyrus, SOG represents the superior occipital gyrus, and STG represents the superior temporal gyrus.

Meta-analytic heterogeneity was low for both clusters (*I² < 39%*), indicating consistent patterns of activation across studies. Jackknife sensitivity analysis demonstrated that all significant clusters remained robust when each study was systematically removed in turn (22/22 studies preserved). Egger’s test showed no evidence of publication bias (*p > 0.05*), supported by visual inspection of funnel plots (see *Supplementary Figure 1*).

These findings provide meta-analytic evidence for convergent recruitment of right-lateralized temporo-parietal regions in visual experts, suggesting the involvement of integrative visual-semantic functions that generalize across domains.

### 3.3. Subgroup Analysis Results

#### 3.3.1. Stimulus-type Subgroup Results

Stimulus-type subgroup analyses revealed distinct lateralization patterns associated with symbolic versus pictorial expertise.

For symbolic stimuli (e.g., musical notation, written language), a left-lateralized temporal cluster was identified, encompassing the left MTG, left superior temporal gyrus (STG), and left inferior temporal gyrus (ITG). The peak activation was located in the left ITG (*MNI: –62, –56, –10; SDM-Z = 3.281; p_TFCE-corr_ <0.05; cluster size = 1207 voxels*). These regions are associated with lexical-syntactic and semantic processing, forming a language-related profile distinct from the core network observed in the primary analysis. An additional activation cluster was detected in the right AG and right MOG, with a local peak in the right MOG (*MNI: 36, –66, 38; SDM-Z = 3.958; p_TFCE-corr_ <0.05; cluster size = 218 voxels*), suggesting a partial overlap with domain-general visual-semantic networks. See *Table 2 and Figure 3A* for details.

**Figure 3.**
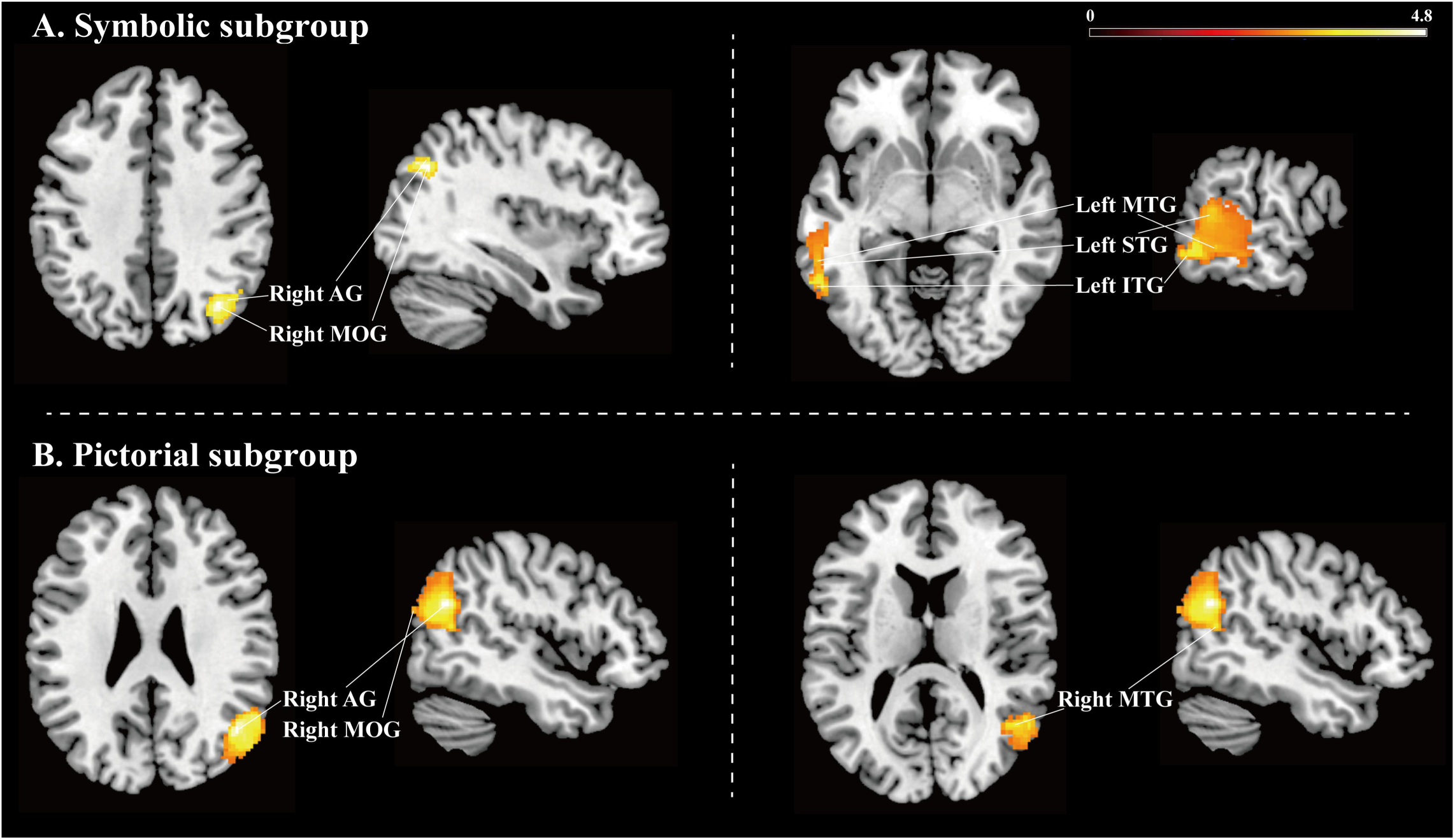
Z-map of significant activations results associated with the visual expertise effect identified by the symbolic and pictorial subgroups meta-analysis at *p_TFCE-corr_* < 0.05. *Note: In heat map representations, a stronger yellow indicates a higher positive z value.

For pictorial stimuli (e.g., radiographs, chessboards), a right-lateralized parieto-temporo-occipital cluster was identified, closely overlapping with the primary meta-analytic results. Robust activation was found in the right AG, right MTG and MOG, with peak activation located in the right AG (*MNI: 46, –62, 26; SDM-Z = 4.581; p_TFCE-corr_ <0.05; cluster size = 1118 voxels*). This pattern reflects the predominant contribution of pictorial domains to the overall activation observed in the full sample. See *Table 2 and Figure 3B* for details.

Meta-analytic heterogeneity estimates were low for both subgroups (*I² < 39%*). Jackknife sensitivity analysis revealed limited stability for the symbolic cluster (4/8 studies preserved), suggesting that these findings may be influenced by study-level variability. In contrast, the pictorial cluster showed strong replicability (13/14 studies preserved). Egger’s tests indicated no significant publication bias in either subgroup (*p > 0.05*; *see Supplementary Figure 1*).

These findings suggest that stimulus type modulates the lateralization of visual expertise-related brain activity, with symbolic domains preferentially engaging left-lateralized language-related temporal regions, and pictorial domains relying on right-lateralized visual-semantic circuits.

#### 3.3.2. Experimental contrast Subgroup Results

Subgroup analyses based on experimental contrast examined whether inter-group (expert vs. novice) versus intra-group (domain-relevant vs. irrelevant stimuli within experts) designs yield distinct neural activation patterns associated with visual expertise.

The inter-group comparison subgroup revealed widespread activation in a bilateral parieto-temporo-occipital circuit (*Table 2; Figure 5*). A large cluster centered in the right AG (*MNI: 54, –56, 26; SDM-Z = 4.474; p_TFCE-corr_ < 0.05; cluster size = 3199 voxels*) extended to the right MTG, right AG, right STG, right MOG, right IPL, right rolandic operculum (RO), right Heschl’s gyrus (HG), and right supramarginal gyrus (SMG). Another cluster peaked in the left fusiform gyrus (FG) (*MNI: –36, –56, –20; SDM-Z = 3.714; p_TFCE-corr_ < 0.05; cluster size = 947 voxels*), including the adjacent left ITG and left inferior occipital gyrus (IOG). A third cluster was identified in the left IFG-triangular (*MNI: –52, 24, 10; SDM-Z = 3.192; p_TFCE-corr_ < 0.05; cluster size = 226 voxels*), with extension into the left insula.

Heterogeneity was very low (*I² <* 39%), indicating high consistency across studies. Jackknife sensitivity analysis demonstrated strong replicability (16/17 studies preserved). Egger’s test and funnel plot inspection (*Supplementary Figure 1*) revealed no significant publication bias (*p > 0.05*).

In contrast, the intra-group comparison subgroup—which compared expert responses to domain-relevant versus irrelevant stimuli—did not yield any significant activation after TFCE correction. This absence of effect may reflect insufficient statistical power given the small number of studies (n = 5), increasing the likelihood of Type II error.

Overall, these findings suggest that experimental contrast may modulate the sensitivity of detecting expertise-related neural activation, with inter-group designs potentially providing a more reliable means of isolating group-level differences in functional engagement.

### 3.4. Meta-regression Analysis Results

Meta-regression analyses were conducted to examine whether continuous demographic variables—male ratio and mean age of expert participants—modulated voxel-wise brain activation patterns associated with visual expertise.

When male ratio was entered as a single regressor, significant lateralized patterns of activation were identified (*Table 2; Figure 6A*), with positive activations in the right MTG, right MOG, and right AG, and negative activations in the left MTG, left STG, left IPL, left AG, and left superior parietal gyrus (SPG). Peak coordinates were located in the right MTG (*MNI: 48, –64, 18; SDM-Z = 4.415; p_TFCE-corr_ < 0.05; cluster size = 576 voxels*), left MTG (*MNI: –62, –32, –4; SDM-Z = –3.066; p_TFCE-corr_ < 0.05; cluster size = 719 voxels*), and left IPL (*MNI: –34, –68, 46; SDM-Z = –2.983; p_TFCE-corr_ < 0.05; cluster size = 442 voxels*).

When both male ratio and mean age were included as covariates, a similar but slightly more spatially restricted pattern was observed (*Table 2; Figure 6B*). Positive activations remained in the right MTG, right MOG, and right AG, while negative activations persisted in the left MTG, left STG, left IPL, and left AG. Peak coordinates were located in the right MTG (*MNI: 52, –68, 18; SDM-Z = 4.675; p_TFCE-corr_ < 0.05; cluster size = 527 voxels*), left MTG (*MNI: –58, –26, –6; SDM-Z = –3.153; p_TFCE-corr_ < 0.05; cluster size = 458 voxels*), and left IPL (*MNI: –34, –68, 46; SDM-Z = –3.027; p_TFCE-corr_ < 0.05; cluster size = 313 voxels*). These results suggest that adding age as a covariate did not meaningfully alter the spatial configuration of expertise-related activations beyond the effects of gender.

Heterogeneity across voxels was generally low (*I²* < 39%), except for the left MTG peak in the gender regression model, which showed moderate variability (40% < *I²* < 59%). Jackknife sensitivity analyses demonstrated high robustness, with all significant clusters preserved in all iterations (22/22 studies). Egger’s tests and visual inspection of funnel plots (*Supplementary Figure 1*) revealed no evidence of publication bias (*p > 0.05*).

These results suggest that, in the current dataset, gender composition showed a more detectable association with lateralization than age.

## 4. Discussion

To evaluate the core–adaptive hypothesis, we pooled 210 peak-activation foci from 22 whole-brain task-fMRI studies that together sampled 579 participants, including 314 recognized experts, across 11 real-world non-face domains. The analysis converged on a right-lateralized parieto-temporo-occipital circuit—centered on the MOG, MTG, AG and neighboring IPL—that consistently recruited across domains and is well positioned to support visuo-semantic integration and attention-guided object recognition. This convergence is consistent with a domain-general “core” scaffold for visual expertise. Subgroup and meta-regression tests indicated that the primary-defined “core” was largely shared across analyses, whereas additional left-hemisphere regions emerged as adaptive extensions contingent on stimulus type, experimental contrast, and gender. Collectively, these results provide convergent cross-domain evidence for a stable right-hemisphere backbone that is flexibly augmented by left-hemisphere systems, unifying disparate domain-specific findings within a single framework and offer a whole-brain reference space for future studies.

### 4.1. A Right-Lateralized “Core” Network Consistently Recruited Across Specialized Domains of Visual Expertise

Our primary meta-analysis (*Table 2, Figures 2, and Figure 6*) provides compelling evidence that visual expertise consistently engages a right-lateralized parieto-temporo-occipital “core” circuit. Specifically, robust activation was observed in the right AG, right MTG, right MOG, and right IPL—regions implicated in visual processing, attentional allocation, and the integration of perceptual inputs with task-relevant semantic representations shaped by experience.

The right AG, located in the posterior inferior parietal lobule, has been implicated as a multimodal convergence hub involved in concept retrieval, semantic integration, and visuospatial attention (Binder et al., 2009; Seghier, 2013). In the context of visual expertise, strong AG activation likely facilitates the mapping of visual features onto semantic representations, thereby supporting expert-level object recognition. The right MTG, specifically its posterior portion as revealed by our activation cluster (Xu et al., 2015), lies along the early stages of the ventral visual stream and is implicated in visual feature processing during object recognition (Bilalić, 2017). Prior work by Bilalić and colleagues has reported this right posterior MTG to differentiate experts from novices in domains such as chess, emphasizing its role in object perception (Bilalić, 2017), in line with the present findings. The right MOG contributes to category-level object perception (Ishai et al., 2000), spatial location encoding (Liu et al., 2017), and the allocation of spatial attention (Cona and Scarpazza, 2019; Hahn et al., 2006). In the context of visual expertise, enhanced activation of the right MOG may reflect domain-specific optimization for processing spatial relationships and salient object features. Notably, the right IPL (excluding the AG and SMG) is part of the dorsal attention network that supports the top-down allocation of attention based on current task goals (Cabeza et al., 2008; Seghier, 2013). Its engagement in visual expertise may reflect the use of goal-directed attentional templates that prioritize task-relevant features during rapid object recognition in complex visual contexts.

Notably, our cross-domain whole-brain synthesis did not yield convergent peaks in the FFA, despite early neuroimaging reports of expertise-related selective activation in the FFA (Burns et al., 2019; Gauthier et al., 1999). Because most FFA–expertise effects are reported from localizer-defined ROIs rather than whole-brain coordinate maps, many such studies were excluded by our inclusion criteria, which likely contributes to the lack of convergent FFA peaks (Burns et al., 2019; Gauthier et al., 1999; Kanwisher et al., 1997). This discrepancy can be interpreted in two ways. First, effects detectable within a pre-specified ROI may not survive stringent whole-brain multiple-comparison control when the same contrast is tested voxelwise across the brain (Nichols and Hayasaka, 2003; Poldrack, 2007). Second, FFA involvement may be expressed less as a consistent univariate peak and more as distributed representational patterns, whereas coordinate-based SDM-style meta-analyses reconstruct maps from reported peak locations and their associated statistics and therefore do not directly test multiregional pattern information (Albajes-Eizagirre et al., 2019). Therefore, the present meta-analysis is largely insensitive to localizer-defined or multivariate FFA expertise effects, and the absence of convergent FFA peaks should not be taken as evidence of absence. Accordingly, the present findings should be viewed as complementary to, rather than competitive with, fusiform-centered accounts of visual expertise.

In summary, by synthesizing whole-brain activation data across diverse specialized visual expertise domains, the present meta-analysis reveals a consistently recruited right-lateralized parieto-temporo-occipital circuit. This convergence underscores the presence of a domain-spanning neural core that integrates perceptual, semantic, and attentional processes to support expert-level visual recognition. These findings highlight that, despite differences in domain-specific tasks and methodological or demographic heterogeneity, visual expertise consistently engages a shared neural scaffold—supporting the “core” component of the core–adaptive hypothesis.

### 4.2. Flexible Left-Lateralized “Adaptive” Systems Modulated by Heterogeneous Factors

Across subgroup and meta-regression analyses, primary-defined right-lateralized parieto-temporo-occipital “core” nodes were largely shared across conditions (*Figure 6; grid-filled labels*), while left-lateralized regions (MTG/STG/ITG/IFG; plus, FG/IOG/insula under higher demands) are flexibly recruited as adaptive extensions depending on stimulus format, contrast type, and sample composition.

#### 4.2.1. Stimulus Type Modulates the Lateralization of Visual Expertise

Within this core–adaptive organization, subgroup analyses suggested that stimulus format—symbolic versus pictorial—elicits distinct adaptive extensions within the core–adaptive framework (*Table 2, Figures 3, and Figure 6*).

Symbolic domains (e.g., musical notation, written language) exhibited a left-lateralized temporal extension to the right-lateralized core network, engaging regions including the left MTG, STG, and ITG—regions preferentially recruited when visual recognition requires rule-based decoding and symbolic-semantic mapping, in contrast to the right-lateralized visual-semantic integration that characterized the primary meta-analysis findings. The left MTG supports the retrieval of lexical–syntactic information from long-term memory (Snijders et al., 2010), facilitating the interpretation of abstract rule-based symbols. The left STG contributes to phonological processing and speech comprehension (Yi et al., 2019), suggesting possible covert verbal rehearsal during symbolic recognition. The left ITG contributes to semantic integration and conceptual access (Price, 2012), supporting abstract symbol comprehension. Notably, symbolic expertise also engaged right-hemisphere components of the core network, including the AG and MOG. This coactivation supports a recruitment pattern in which symbolic recognition weights left-lateralized rule-based decoding while preserving engagement of the right-lateralized visual-semantic core. Importantly, we do not equate adaptive recruitment with verbal strategy. Rather, language-related regions likely constitute one instantiation of adaptive extension when representational formats favor symbolic decoding.

In contrast, pictorial object expertise (e.g., radiographs, chessboards) consistently activated right MTG, AG, and MOG—regions largely consistent with the core network identified in the primary meta-analysis—thereby providing further evidence for the robustness and domain-generality of the right-lateralized parieto-temporo-occipital circuit. This convergence reflects the predominant contribution of pictorial domains to the overall activation observed in the full sample. The right-lateralized activation indicates an engagement of visual-semantic integration systems optimized for recognizing complex, visually grounded objects.

Collectively, these findings indicate that visual expertise is not instantiated by a fixed neural signature, but flexibly adapts to stimulus type. Symbolic expertise partially recruits the right-lateralized core network along with a left temporal extension supporting syntactic decoding (Friederici, 2011), whereas pictorial expertise primarily engages the core network for visual-semantic integration optimized at the feature level (Bukach et al., 2006; Grill-Spector and Weiner, 2014). These patterns suggest that stimulus type-specific idiosyncrasies modulate the weighting of shared components and recruit additional circuitry as needed, illustrating the flexible, content-sensitive nature of the adaptive system.

#### 4.2.2. Experimental Contrast Influences the Breadth of Adaptive Recruitment Beyond the Core

Beyond intrinsic task features, experimental design—particularly inter-versus intra-group comparisons—shaped the spatial extent of adaptive recruitment, revealing design-sensitive expansions beyond the core network (*Table 2, Figures 4, and Figure 6*).

**Figure 4.**
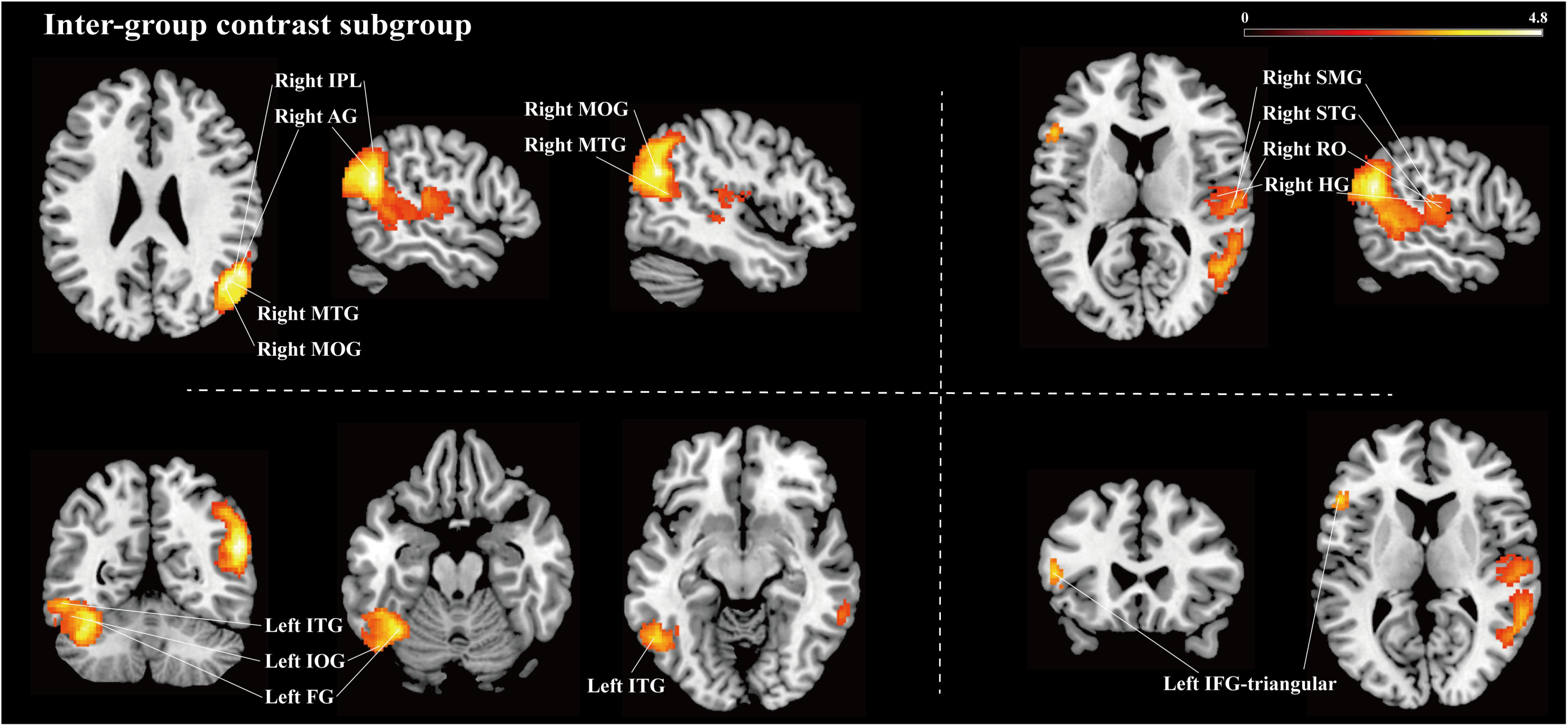
Z-map of significant activations results associated with the visual expertise effect identified by the inter-group comparison subgroup meta-analysis at *p_TFCE-corr_* < 0.05. *Note: In heat map representations, a stronger yellow indicates a higher positive z value.

Intra-group comparison subgroup—contrasting expert responses to domain-relevant versus irrelevant stimuli—did not yield statistically significant clusters, likely due to insufficient power from the limited number of available studies (n = 5). However, it may also reflect the fact that, within experts, the cognitive distinction between domain-relevant and irrelevant stimuli does not elicit sufficiently robust or consistent neural differentiation to be detected at the meta-analytic level. This suggests that intra-group contrasts, especially with current sample sizes, may be less likely to yield consistent whole-brain convergence at the current scale of the literature, compared with inter-group designs.

By contrast, the inter-group contrast subgroup (e.g., experts vs. novices) revealed an expanded bilateral network with right-hemisphere predominance, not only re-engaging core regions of the right parieto-temporo-occipital circuit (AG, MTG, MOG, IPL), but also recruiting additional bilateral areas—including right STG, RO, HG, SMG, as well as left ITG, FG, IOG, IFG-triangular, and insula. The right STG—minimally activated in the primary analysis—exhibited reliable recruitment here, implicated in phonological processing and speech comprehension, it may support expert object recognition via covert verbal labeling (Yi et al., 2019). The right RO, involved in sensorimotor integration, may facilitate internal verbal rehearsal or action simulation during expert visual recognition (Koelsch et al., 2006). The right HG, situated in primary auditory cortex, may support covert verbal encoding of diagnostic visual features during expert object recognition (Warrier et al., 2009). The right SMG, implicated in phonological decision-making, may contribute to expert-level categorization through cross-modal verbal support (Hartwigsen et al., 2010). The left ITG supports semantic integration of complex visual inputs (Price, 2012); the left FG is critical for rapid and efficient visual processing of written language (McCandliss et al., 2003); the left IOG contributes to early-stage visual feature extraction (Sato et al., 2016); the left IFG-triangular is critical for semantic selection and integration (Snijders et al., 2010; Thompson-Schill et al., 1997). The left insula, identified as a core region in language processing—including comprehension, production, and lexical learning—may support visual expertise by facilitating semantic integration and learning during object recognition (Ardila et al., 2014).

Together, these findings suggest that experimental contrast modulates the breadth of adaptive engagement beyond the right-lateralized core. Inter-group designs—by maximizing visual and semantic differentiation—elicit expanded bilateral recruitment involving verbal, sensorimotor, and semantic systems. This contrast-driven modulation not only affirms the robustness of the core network but also highlights how inter-group comparisons dynamically extend the recruitment of adaptive systems.

#### 4.2.3. Demographic Factors Influence Lateralization of Visual Expertise

Meta-regression analyses revealed that demographic factors, particularly gender composition, influenced the hemispheric lateralization of adaptive recruitment within the visual expertise network (*Table 2, Figures 5, and Figure 6*).

**Figure 5.**
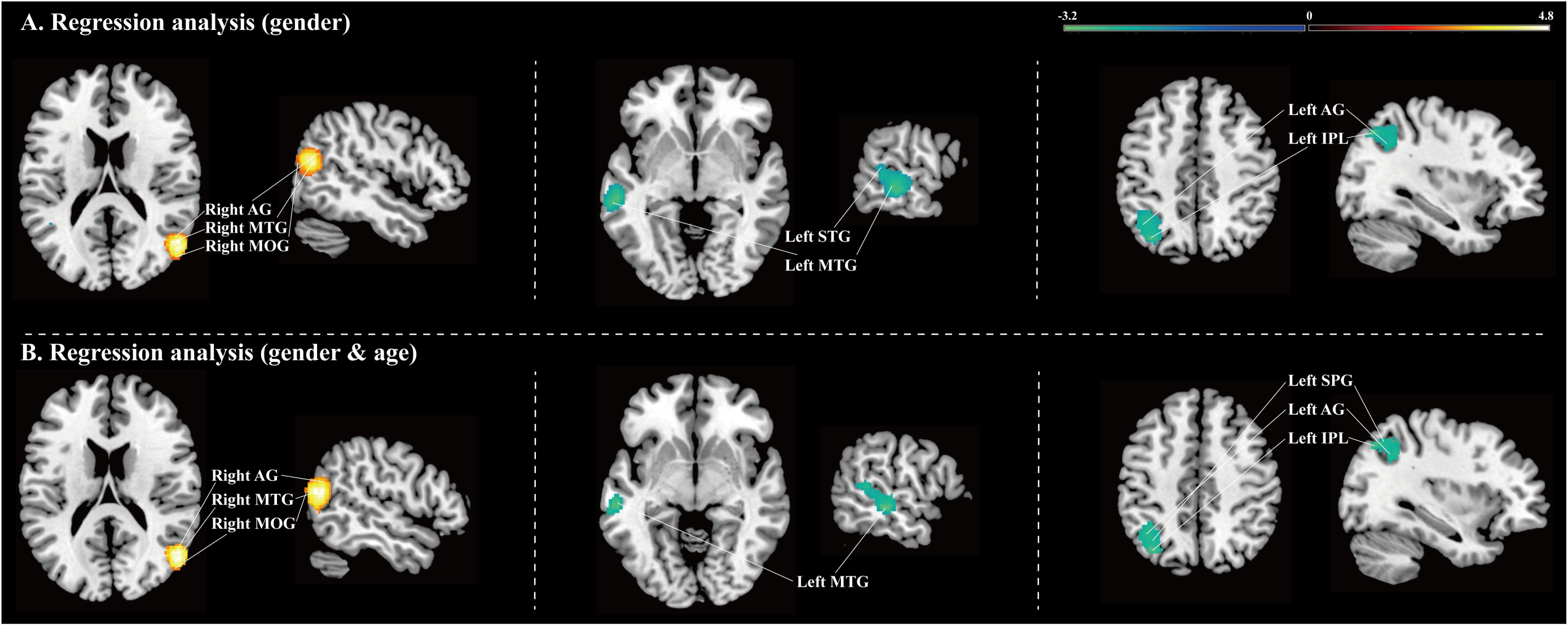
Z-map of significant activations results associated with the visual expertise effect identified by the meta-regression analysis at *p_TFCE-corr_* < 0.05. *Note: In heat map representations, a stronger yellow indicates a higher positive z value, and a brighter green represents a larger negative z value.

**Figure 6.**
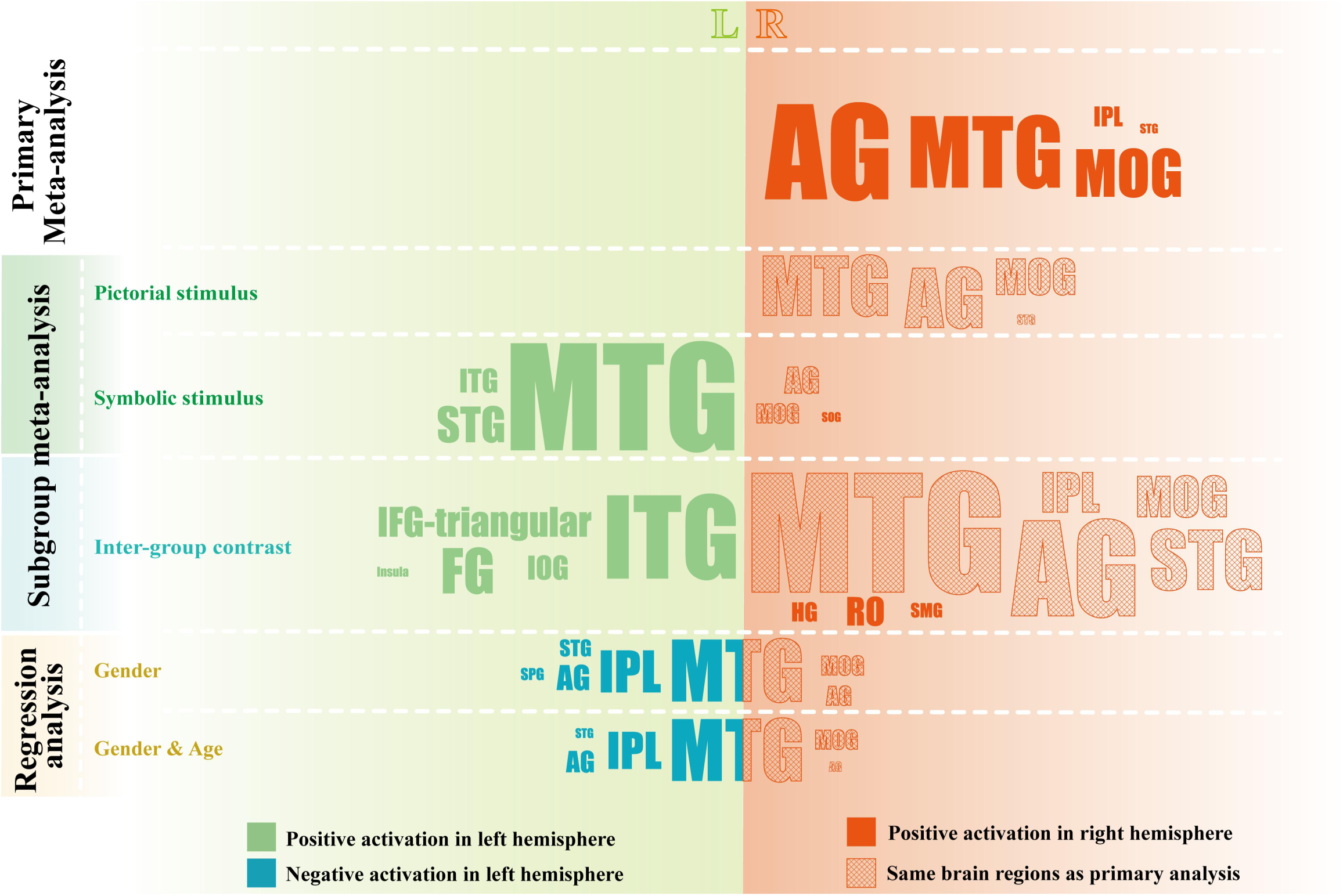
Word-cloud summary of the core–adaptive architecture across analyses. Word size is proportional to cluster extent (larger = more voxels). The background is divided into two colour fields: an orange-red tint on the right marks the right hemisphere and a light-green tint on the left marks the left hemisphere. Brain region names printed in the matching orange-red or light-green type indicate significant positive activations, while blue type marks significant negative activations (*p_TFCE-corr_ <0.05*). Grid-filled brain-region labels denote primary-defined “core” nodes (i.e., clusters identified in the primary meta-analysis), whereas non–grid-filled labels indicate additional clusters that emerge only in specific subgroup or regression analyses (adaptive extensions).

After controlling for male proportion (71.05% across studies), robust activation emerged in core right-hemisphere regions, specifically the right MTG, MOG, and AG, thereby replicating the primary meta-analysis and highlighting the stability of the right-lateralized core network across demographic subgroups. In contrast, decreased activation was observed in left-hemisphere regions, including the left MTG, STG, IPL, AG, and SPG—all canonically involved in language-related processing. The left MTG supports lexical–syntactic retrieval (Snijders et al., 2010); the left STG contributes to auditory-verbal integration (Yi et al., 2019); the left IPL is associated with semantic comprehension (Numssen et al., 2021); the left AG plays a broad role in concept retrieval and semantic integration (Seghier, 2013); and the left SPG involves in linguistic writing processes (Magrassi et al., 2010). Notably, none of these left-lateralized suppressions appeared in the primary meta-analysis, indicating that the male-skewed sample may have masked the detection of left-hemisphere deactivation associated with visual expertise. This pattern suggests that gender composition may act as a moderator of hemispheric asymmetry at the group level, rather than indicating sex-specific mechanisms of expertise.

When age was added as a covariate, the activation pattern remained largely unchanged, except for the absence of suppression in the left SPG. This minimal shift indicates that, in the available literature, lateralization appears more sensitive to gender composition than to age-related variability (noting male-skewed samples). One possible explanation is that age-related effects overlap with expertise acquisition due to shared mechanisms of long-term training-induced plasticity; in addition, the relatively narrow age range across studies (e.g., 14 out of 22 studies had a mean age between 21 and 30) may have limited the ability to detect independent age-related variability, further amplifying the observed gender-driven effects.

Together, these findings support that while the right-lateralized core network remains consistently engaged across demographic subgroups, adaptive engagement of left-hemisphere regions varies by gender composition—reflecting the demographic sensitivity of the core–adaptive architecture. This underscores the importance of considering sample diversity when interpreting neural signatures of visual expertise.

### 4.3. Non-Face Visual Expertise Recruits a Distinct Whole-Brain Architecture from Face Processing

Faces and non-face expertise place partly different demands on the cognitive operations that support skilled recognition. Face recognition is intrinsically embedded in social communication: beyond extracting visual structure, observers routinely infer identity, person knowledge, and affective meaning, such that even ostensibly “perceptual” face tasks can invite socially meaningful interpretation (Haxby et al., 2000). Moreover, face processing is shaped by early developmental constraints, with evidence consistent with a sensitive-period contribution to typical holistic/configural face perception (McKone et al., 2007). In contrast, non-face real-world expertise is typically built through prolonged, deliberate practice beyond early infancy, and is more plausibly expressed as experience-driven refinement within mature cognitive systems—prioritizing high-precision discrimination, efficient visual–semantic mapping, and goal-directed attentional selection under task constraints.

These differences at the level of cognitive demands are mirrored in the neuroimaging literature. Whole-brain ALE meta-analyses of face processing converge on a distributed system anchored in occipito-temporal face-responsive regions (IOG/FFG and pSTS/MTG) and extending to limbic and frontal components that support affective–social significance and task-dependent evaluative readout (e.g., amygdala, IFG/pre-SMA) (Müller et al., 2018b). When tasks explicitly require affective decoding, convergence further emphasizes an amygdala–STS–frontal/medial control configuration (Dricu and Frühholz, 2016), whereas social-evaluative judgments (e.g., trustworthiness/attractiveness) additionally implicate valuation-related circuitry such as OFC/ventral striatum (Bzdok et al., 2011). Against this face-processing profile, our cross-domain meta-analysis identifies a convergent whole-brain signature of non-face visual expertise centered on a right temporo-parietal–occipital network. Nevertheless, taken together, these literatures—juxtaposed here as a theoretical rather than statistical comparison, rather than as a direct voxelwise contrast within a single meta-analytic framework—are consistent with the view that non-face visual expertise and face processing may emphasize partially distinct cognitive–neural representational regimes. Face processing is more tightly anchored in ventral occipito-temporal visual analyses with reliable affective and social-evaluative extensions, whereas non-face expertise converges on a right-lateralized temporo-parietal–occipital scaffold that supports precise integration and visual–semantic binding across trained domains. These observations motivate future work that directly quantifies similarities and differences between face and non-face expertise.

### 4.4. Limitations

Several limitations should be considered when interpreting the present findings. First, coordinate-based meta-analysis relies on peak activation coordinates rather than full statistical images, reducing spatial precision and preventing direct estimation of effect-size magnitudes within clusters. Second, although moderator analyses mitigated some variability, the contributing studies remain heterogeneous in expert-definition standards, training background, behavioral baselines, and analysis pipelines; unmeasured factors (e.g., expertise depth, training duration, and strategy) may still confound the results. Third, publication bias cannot be ruled out because null or weak-effect studies are under-represented in the neuroimaging literature and, by extension, in our database. Fourth, subgroups of interest—such as intra-group contrasts or symbolic stimuli—contained fewer than ten studies, limiting statistical power, jackknife stability and detection of subtle moderator effects. As the literature grows, larger and more balanced datasets will permit finer-grained analyses of task, domain, and demographic interactions. Fifth, because our synthesis is limited to non-face expertise and relies on reported peaks, we could not perform a direct, quantitative face–non-face comparison within a single meta-analytic space; future work should explicitly quantify similarities and differences across the two literatures. Finally, all the studies included employed cross-sectional designs; longitudinal or training studies are needed to establish causal links between neural plasticity and expertise acquisition.

### 4.5. Conclusion

By synthesizing whole-brain task-based fMRI data from 22 studies spanning 11 distinct types of non-face expertise, this meta-analysis delineates a core–adaptive neural architecture for real-world non-face visual expertise. A right-lateralized parieto-temporo-occipital core integrates fine-grained visual features with semantic knowledge and supports attention-guided object recognition, while left-hemisphere adaptive systems are recruited according to stimulus format, contrast design, and demographic context. These findings extend current theories of perceptual expertise and offer a domain-spanning whole-brain reference framework for future studies.

## Declaration of competing interests

The authors declare no competing interests.

## Acknowledgments

This paper is supported by the National Key R&D Program of China (Grant No.2022YFF1202400).

## Declaration of generative AI and AI-assisted technologies in the writing process

During the preparation of this work the author(s) used a large language model (LLM) [GPT-4/ OpenAI] in order to solely refine linguistic aspects and improve readability. After using this LLM, the authors reviewed and edited the content as needed and take full responsibility for the content of the published article. Any perspectives or interpretations expressed are exclusively those of the authors and do not represent the views of the LLM provider or its affiliates.

## Data availability

The data supporting the findings of this study can be accessed on the Open Science Framework (OSF) platform at https://osf.io/dpcy5/files/osfstorage.

## Notes

https://osf.io/dpcy5

https://osf.io/dpcy5/files/osfstorage

### Competing Interest Statement

The authors have declared no competing interest.

