## Supplemental Data 1 for "Whole-Brain Convergence of Real-World Visual Expertise beyond Faces: Meta-Analytic Evidence for Core–Adaptive Neural Architecture"

### Supplementary material

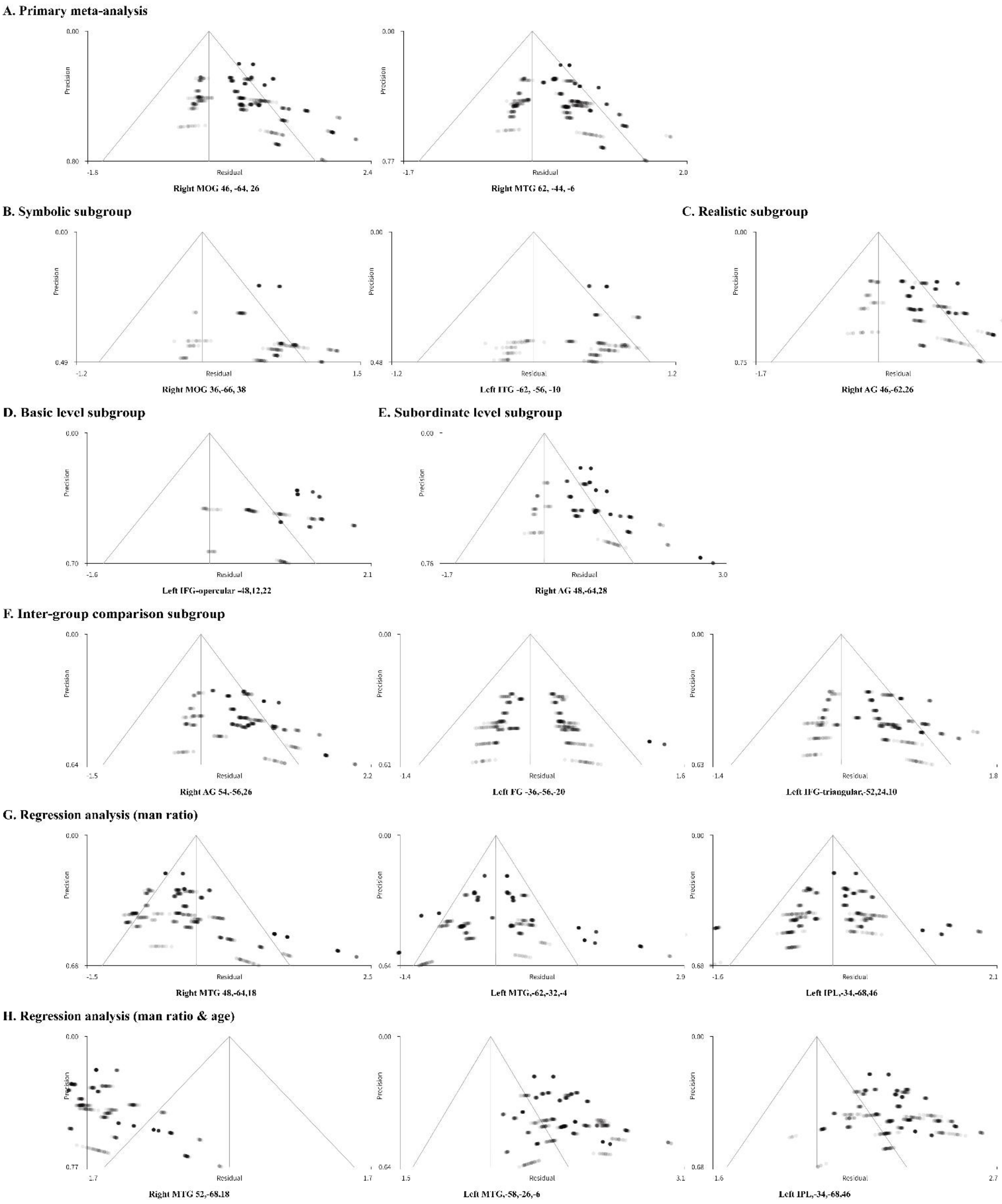


**Supplementary Figure 1 |** Funnel plots for the clusters revealed in the meta-analyses.

Funnel plots are shown for peak coordinates from all significant clusters identified in the primary meta-analysis, subgroup analyses, and meta-regressions.
*Note*: AG represents the angular gyrus, FG represents the fusiform gyrus, IFG-triangular represents the inferior frontal gyrus, triangular part, IPL represents the inferior parietal (excluding supramarginal and angular) gyri, ITG represents the inferior temporal gyrus, MOG represents the middle occipital gyrus, MTG represents the middle temporal gyrus.

**Supplementary Table 1 |** Search formulas in PubMed, PsycINFO, and Web of Science.

| |
|---|
| **PubMed:** |
| ("fMRI"[Text Word]) AND (("expert"[Text Word]) OR "expertise"[Text Word] OR "professional"[Text Word] OR "training"[Text Word] OR "train"[Text Word] OR "prodigy"[Text Word]) AND (("visual"[Text Word]) OR "vision"[Text Word]) NOT (("monkey"[Text Word]) OR "mice"[Text Word] OR "animal"[Text Word] OR "patient"[Text Word]) |
| **PsycINFO:** |
| (fMRI) AND (expert* OR expertise OR professional* OR training OR train OR prodigy) AND (visual OR vision) NOT (monkey OR mice OR animal OR patient) |
| **Web of Science:** |
| TS=(fMRI) AND TS=(expert OR expertise OR professional OR training OR train OR prodigy) AND TS=(visual OR vision) NOT TS=(monkey OR mice OR animal OR patient) |

**Supplementary Table 2 |** Quality assessment checklists (Score 0/0.5/1 for each item, total score 15 out of 15).

| |
|---|
| **Subjects** |
| **Item 1:** Expert participants were evaluated prospectively, using strict assessment criteria to confirm their professional expertise and ensure they were in good health, with demographic data provided.<br>**Item 2:** Non-expert control participants were evaluated prospectively, with strict criteria to confirm the absence of relevant expertise and ensure they were in good health, along with demographic data provided.<br>**Item 3:** Key variables such as age, gender, and handedness were accounted for, either through stratification or statistical methods.<br>**Item 4:** Withdrawals from the study were explained (scores 1), or not (scores 0.5).<br>**Item 5:** Group sample size: ≥ 20 participants, scores 1; > 10 participants, scores 0.5. |
| **Method for tasks** |
| **Item 6:** All participants were required to perform tasks during the scan that specifically involved visual recognition processes.<br>**Item 7:** Studies must include explicit comparisons of visual expertise effects: inter-group studies comparing visual experts with non-experts on tasks relevant to their expertise, or intra-group studies comparing tasks related to expertise with tasks unrelated to expertise among visual experts. |
| **Method for image acquisition and statistical analysis** |
| **Item 8:** The imaging technique used was described in sufficient detail for replication.<br>**Item 9:** Appropriate analytical methods were used, with explicit mention of whole-brain analysis (score 1) or inferred from the study context that whole-brain analysis was used (score 0.5).<br>**Item 10:** Statistical methods were suitably applied for the primary effect analysis.<br>**Item 11:** Corrections for multiple statistical comparisons were performed using whole-brain correction (e.g., FWE, FDR, cluster-level; score = 1), ROI-based correction (score = 0.5), or not corrected (score |

= 0).

| Results, conclusions and conflict of interest |
|---|
| **Item 12:** Statistical values for both significant and non-significant differences were reported. |
| **Item 13:** Spatial coordinates were reported in a standard space (e.g., MNI or Talairach coordinates). |
| **Item 14:** Conclusions were consistent with the results obtained and the limitations were discussed. |
| **Item 15:** Declarations of conflict of interest or identification of finding sources |

**Supplementary Table 3 |** Quality assessment scores of included papers

| Included papers | Items | | | | | | | | | | | | | | | Quality scores |
|---|---|---|---|---|---|---|---|---|---|---|---|---|---|---|---|---|
| | Subjects | | | | | Method for tasks | | Method for image acquisition and statistical analysis | | | | Results, conclusions and conflict of interest | | | | |
| | 1 | 2 | 3 | 4 | 5 | 6 | 7 | 8 | 9 | 10 | 11 | 12 | 13 | 14 | 15 | |
| Daniele Schön et al. [1] | 1 | 1 | 0.5 | 0.5 | 0 | 1 | 1 | 1 | 0.5 | 1 | 1 | 1 | 1 | 0.5 | 1 | 12 |
| Geng Li et al. [2] | 1 | 1 | 1 | 0.5 | 0.5 | 1 | 1 | 1 | 0.5 | 1 | 1 | 1 | 1 | 0.5 | 1 | 13 |
| Campitelli et al. [3] | 1 | 1 | 1 | 1 | 0 | 1 | 1 | 1 | 0.5 | 1 | 0.5 | 1 | 1 | 0.5 | 1 | 13 |
| Bilalic et al. [4] | 1 | 1 | 1 | 0.5 | 0.5 | 1 | 1 | 1 | 1 | 1 | 1 | 1 | 1 | 0.5 | 1 | 13.5 |
| Lee et al. [5] | 1 | 1 | 1 | 0.5 | 0 | 1 | 1 | 1 | 0.5 | 1 | 0.5 | 1 | 1 | 0.5 | 1 | 12.5 |
| Hoenig et al. [6] | 1 | 1 | 1 | 0.5 | 1 | 1 | 1 | 1 | 0.5 | 1 | 1 | 1 | 1 | 0.5 | 1 | 13.5 |
| Marcin Szwedet al. [7] | 1 | 1 | 1 | 0.5 | 0.5 | 1 | 1 | 1 | 1 | 1 | 1 | 1 | 1 | 1 | 1 | 14 |
| Valeria Mongelli et al. [8] | 1 | 1 | 1 | 1 | 1 | 1 | 1 | 1 | 1 | 1 | 1 | 1 | 1 | 1 | 1 | 15 |
| Farah Martens et al. [9] | 1 | 1 | 1 | 1 | 1 | 1 | 1 | 1 | 1 | 1 | 1 | 1 | 1 | 1 | 1 | 15 |
| Wanwan Guo et al. [10] | 1 | 1 | 1 | 1 | 1 | 1 | 1 | 1 | 0 | 1 | 1 | 1 | 1 | 1 | 1 | 15 |
| Wong et al. [11] | 1 | 1 | 1 | 1 | 0.5 | 1 | 1 | 1 | 0.5 | 1 | 1 | 1 | 1 | 0.5 | 1 | 13.5 |
| Krawczyk et al. [12] | 1 | 1 | 1 | 0.5 | 0 | 1 | 1 | 1 | 0.5 | 1 | 0 | 1 | 1 | 1 | 1 | 12.5 |
| Bilalić et al. [13] | 1 | 1 | 1 | 0.5 | 0 | 1 | 1 | 1 | 0.5 | 1 | 0.5 | 1 | 1 | 0.5 | 1 | 12.5 |
| Wu et al. [14] | 1 | 1 | 1 | 1 | 0.5 | 1 | 1 | 1 | 1 | 1 | 0.5 | 1 | 1 | 0.5 | 1 | 14 |
| Bilalić et al. [15] | 1 | 1 | 1 | 0.5 | 0.5 | 1 | 1 | 1 | 1 | 1 | 1 | 1 | 1 | 0.5 | 1 | 13.5 |
| Bartlett et al. [16] | 1 | 1 | 1 | 0.5 | 0.5 | 1 | 1 | 1 | 1 | 1 | 1 | 1 | 1 | 0.5 | 1 | 13.5 |
| Sharon et al. [17] | 1 | 1 | 1 | 0.5 | 0.5 | 1 | 1 | 1 | 1 | 1 | 1 | 1 | 1 | 0.5 | 1 | 13.5 |
| Sven et al. [18] | 1 | 1 | 1 | 0.5 | 0.5 | 1 | 1 | 1 | 0.5 | 1 | 1 | 1 | 1 | 1 | 1 | 13.5 |
| Marcio et al. [19] | 1 | 1 | 1 | 1 | 1 | 1 | 1 | 1 | 0.5 | 1 | 1 | 1 | 1 | 1 | 1 | 14.5 |
| Duan et al. [20] | 1 | 1 | 1 | 0.5 | 0.5 | 1 | 1 | 1 | 1 | 1 | 1 | 1 | 1 | 0.5 | 1 | 13.5 |

**Supplementary Table 4 |** The PRISMA checklist 2020

| Section and topic | Item # | Checklist item | Reported on page # |
|---|---|---|---|
| **TITLE** | | | |
| Title | 1 | Identify the report as a systematic review. | 1 |
| **ABSTRACT** | | | |
| Abstract | 2 | See the PRISMA 2020 for Abstracts checklist | 2 |
| **INTRODUCTION** | | | |
| Rationale | 3 | Describe the rationale for the review in the context of existing knowledge. | 3 |
| Objectives | 4 | Provide an explicit statement of the objective(s) or question(s) the review addresses. | 3 |
| **METHODS** | | | |
| Eligibility criteria | 5 | Specify the inclusion and exclusion criteria for the review and how meta-analyses were grouped for the synthesis. | 4,5 |
| Information sources | 6 | Specify all databases, registers, websites, organisations, reference lists and other sources searched or consulted to identify studies. Specify the date when each source was last searched or consulted. | 4 |
| Search strategy | 7 | Present the full search strategies for all databases, registers and websites, including any filters and limits used. | 4 |
| Selection process | 8 | Specify the methods used to decide whether a meta-analysis met the inclusion criteria of the review, including how many reviewers screened each record and each report retrieved, whether they worked independently, and if applicable, details of automation tools used in the process. | 5 |
| Data collection process | 9 | Specify the methods used to collect data from reports, including how many reviewers collected data from each report, whether they worked independently, any processes for obtaining or confirming data from study investigators, and if applicable, details of automation tools used in the process. | 5 |
| Data items | 10a | List and define all outcomes for which data were sought. Specify whether all results that were compatible with each outcome domain in each study were sought (e.g., for all measures, time points, analyses), and if not, the methods used to decide which results to collect. | 5 |

| | | | |
|---|---|---|---|
| | 10b | List and define all other variables for which data were sought (e.g., participant and intervention characteristics, funding sources). Describe any assumptions made about any missing or unclear information. | Table 1 |
| Meta-analysis quality assessment | 11 | Specify the methods used to assess quality in the included meta-analyses, including details of the tool(s) used, how many reviewers assessed each meta-analysis and whether they worked independently, and if applicable, details of automation tools used in the process. | 6 |
| Effect measures | 12 | Specify for each outcome the effect measure(s) (e.g., risk ratio, mean difference) used in the synthesis or presentation of results. | 5,6 |
| Synthesis methods | 13a | Describe the processes used to decide which meta-analyses were eligible for each synthesis (e.g., tabulating the study intervention characteristics and comparing against the planned groups for each synthesis. | 5,6,Table 1 |
| | 13b | Describe any methods required to prepare the data for presentation or synthesis, such as handling of missing summary statistics, or data conversions. | 5 |
| | 13c | Describe any methods used to tabulate or visually display results of individual meta-analyses. | 5 |
| | 13d | Describe any methods used to synthesise results and provide a rationale for the choice(s). | 5 |
| | 13e | Describe any methods used to explore possible causes of heterogeneity among study results (e.g., subgroup analysis, meta-regression). | 6 |
| | 13f | Describe any sensitivity analyses conducted to assess robustness of the synthesised results. | 6 |
| Reporting bias assessment | 14 | Describe any methods used to assess risk of bias due to missing results in a synthesis (arising from reporting biases). | 6 |
| Certainty assessment | 15 | Describe any methods used to assess certainty (or confidence) in the body of evidence for an outcome. | - |
| **RESULTS** | | | |
| Study selection | 16a | Describe the results of the search and selection process, from the number of records identified in the search to the number of studies included in the review, ideally using a flow diagram. | 4,5, Figure 1 |
| | 16b | Cite studies that might appear to meet the inclusion criteria, but which were excluded, and explain why they were excluded. | 4,5, Figure 1 |

| | | | |
|---|---|---|---|
| Study characteristics | 17 | Cite each included study and present its characteristics. | 6,7, Table 1 |
| Quality of meta-analyses | 18 | Present assessments of quality for each included meta-analysis. | 7, Supplementary Tables 2 and 3 |
| Results of individual meta-analyses | 19 | For all outcomes, present, for each meta-analysis: (a) summary statistics for each group (where appropriate) and (b) an effect estimates and its precision (e.g., confidence/credible interval), ideally using structured tables or plots. | 7-9, Table 1 and Table 2 |
| Results of syntheses | 20a | For each synthesis, briefly summarise the characteristics and quality among contributing meta-analyses. | 7-9, Figure 2-7 and Table 2 |
| | 20b | Present results of all statistical syntheses conducted. If meta-analysis was done, present for each the summary estimate and its precision (e.g., confidence/credible interval) and measures of statistical heterogeneity. If comparing groups, describe the direction of the effect. | 7-9, Figure 2-7 and Table 2 |
| | 20c | Present results of all investigations of possible causes of heterogeneity among study results. | 7-9, Table 2 |
| | 20d | Present results of all sensitivity analyses conducted to assess the robustness of the synthesised results. | 7-9, Table 2 |
| Reporting biases | 21 | Present assessments of risk of bias due to missing results (arising from reporting biases) for each synthesis assessed. | 7-9, Table 2 |
| Certainty of evidence | 22 | Present assessments of certainty (or confidence) in the body of evidence for each outcome assessed. | - |
| **DISCUSSION** | | | |
| Discussion | 23a | Provide a general interpretation of the results in the context of other evidence | 9-15 |
| | 23b | Discuss any limitations of the evidence included in the review. | 14 |
| | 23c | Discuss any limitations of the review processes used. | 14 |
| | 23d | Discuss implications of the results for practice, policy, and future research | 14 |
| **OTHER INFORMATION** | | | |
| Registration and protocol | 24a | Provide registration information for the review, including register name and registration number, or state that the review was not registered. | 4 |
| | 24b | Indicate where the review protocol can be accessed, or state that a protocol was not prepared. | 4 |

| | | | |
|---|---|---|---|
| | 24c | Describe and explain any amendments to information provided at registration or in the protocol. | - |
| Support | 25 | Describe sources of financial or non-financial support for the review, and the role of the funders or sponsors in the review. | 15 |
| Competing interests | 26 | Declare any competing interests of review authors. | 15 |
| Availability of data, code, and other materials | 27 | Report which of the following are publicly available and where they can be found: template data collection forms; data extracted from included studies; data used for all analyses; analytic code; any other materials used in the review. | 15 |

*From*: Page MJ et al., The PRISMA 2020 statement: an updated guideline for reporting systematic reviews. BMJ 2021;372. doi: 10.1136/bmj.n71. PRISMA 2020 has been originally designed for systematic reviews of individual studies.
